# Cybernetic control of a natural microbial co-culture

**DOI:** 10.1101/2024.07.04.602068

**Authors:** Ting An Lee, Jan Morlock, John Allan, Harrison Steel

## Abstract

A key obstacle in the widespread application of microbial co-cultures in bioprocesses is their compositional instability, as faster-growing species outcompete and dominate the culture. While several synthetic biology approaches have demonstrated control over co-culture composition, there has been an increased interest in computer-based cybernetic control approaches that can offload burdensome genetic control circuitry to computers and enable dynamic control and real-time noise rejection. This work extends that approach, demonstrating a cybernetic control method that is not reliant on any genetic engineering, instead interfacing cells with computers by exploiting their natural characteristics to measure and actuate the composition. We apply this to a *Pseudomonas putida* (*P. putida*) and *Escherichia coli* (*E. coli*) co-culture grown in Chi.Bio bioreactors, first showing how composition estimates calculated from different bioreactor measurements can be combined with a system model using an extended Kalman filter to generate accurate estimates of a noisy system. We also demonstrate that because the species have different optimal temperature niches, adjusting the temperature of the culture can drive the composition in either direction. By using a proportional-integral control algorithm to calculate the temperature that would bring the measured composition towards the desired composition, we are able to track dynamic references and stabilised the co-culture for 7 days (*∼*250 generations), with the experiment ending before the cells could adapt out of the control. This cybernetic framework is broadly applicable, with different microbes’ unique features and specific growth niches enabling robust control over diverse co-cultures.

## Introduction

The use of microbial co-cultures for biotechnological applications has many potential advantages over monocultures: improved productivity from specialisation or splitting of metabolic pathways between species (division of labour) [1, 2], the induction of valuable metabolites [3], increased robustness to contamination [4], or the ability to utilise cheaper substrates by combining strains with different metabolic capabilities [5]. A key obstacle preventing the widespread adoption of co-culturing methods is the challenge posed by engineering and controlling their composition. In particular - and as stated by the competitive exclusion principle [6]- when multiple species compete for a limiting resource in a well-mixed co-culture, the faster-growing species will out-compete and eventually take over, reverting the system to a monoculture. Therefore, a major open challenge in the field is developing robust methods to control and stabilise co-culture composition. Such methods would enable broad applications of co-cultures, and also accelerate their study and exploitation *in general* : the ability to dynamically control co-cultures allows a scientist to quickly explore the composition parameter space and thus maximise bioprocess productivity [7, 8] or probe interspecies dynamics.

To address this control challenge, many engineered biological control systems have been developed to regulate co-culture composition. This includes methods based on complementary auxotrophies [9], quorum sensing circuits [10], burden growth rate actuators [11], and more [12]. While such methods have proven effective for some applications, they have a number of limitations. One is that methods which require genetic engineering of constituent microbes are difficult to implement in complex communities that include non-model organisms and may encounter regulatory challenges if a co-culture is deployed in applications beyond the laboratory. Second is the metabolic burden imparted by genetically encoded parts, which manifests both as the diversion of cellular resources from bioproduction towards governing composition - potentially decreasing productivity - as well as an increased selective pressure against the controller, encouraging escape mutations that break the control circuitry. Both of these limitations are exacerbated if attempting more advanced control with more complex (and thus more burdensome) genetic circuitry.

To mitigate the above challenges, computer-based cybernetic approaches [13]have been used to control composition by shifting some or all of the composition control functions from biological circuitry to computers [14–16]. These control methods make real-time estimates of co-culture composition, compare this measurement to a desired reference composition, compute a control action predicted to drive composition closer to the reference, and then execute these actions. Strengths of cybernetic approaches include noise rejection, the ability to account for uncertain measurements, and to track dynamic references which benefit processes where the optimal composition changes over time [13, 17]. Past realisations of cybernetic co-culture control systems have used computers to *calculate* control actions. However, these actions have typically been *implemented* by genetic engineering of co-culture members to establish suitable computer-cell interfaces. These may include optical [16] or chemical [18] responsive systems introduced to co-culture members to actuate changes in composition, and always relied on fluorescent reporters that are measured as a proxy for species abundance [14–16, 18]. Such engineered biological interfaces allow (comparatively) straightforward implementation of computer-implemented control algorithms, however, control performance is conditional on the continued functioning of the genetically engineered interfaces. This can lead to undesirable fragility - genetically encoded parts (which are not part of the host cell’s core functionality) are often perturbed by mutations, which can change/eliminate their function and hence the co-culture’s behaviour from the controller’s actions. For example, Gutíerrez et al. (2022) demonstrated excellent control of a two-strain *E. coli* co-culture for 40 hours, but after this point escape mutations led to drift despite the computational control component’s attempt to compensate.

We propose that by exploiting natural microbial characteristics and combining several population-averaged measurements, co-culture composition can be controlled with a cybernetic approach that does not require genetically encoded systems for control, measurement, or actuation of composition. For a given two-strain co-culture (1A), this first requires identifying suitable microbial characteristics that might be used to actuate the growth rates, which can be any environmental niche axis where the two strains do not completely overlap. By altering this environmental condition, we can thereby differentially favour the growth of one strain over the other. Culture composition can be determined through projected growth rates under different conditions, with improved accuracy if the strains have any other measurable characteristic that distinguishes them.

Once identified, such a system can be characterised in monocultures to parameterise the relationship between growth rates (or other measured parameters) and chosen control inputs. After testing this relationship in a co-culture, this can then be used to parameterise a mathematical model and design a computer controller. Finally, these can be combined to implement cybernetic control in a co-culture, and the results used to iteratively improve the model and controller if desired.

In this paper, we realise such an approach by demonstrating long-term dynamic control over the composition of a *P. putida* and *E. coli* co-culture. This is achieved by combining simple control algorithms with actuation via small temperature changes, and estimation of co-culture composition by combining measurements of optical density and fluorescence and their variation in time. We initially parameterise growth and production models for each co-culture member in monoculture experiments, then evaluate and adjust their behaviour in co-cultures. In the next step, we apply the model with minimal adjustments to enable control in a co-culture setting, demonstrating its strong performance in over 25 independent bioreactor co-culture experiments. Key innovations of our work include our model-based approach to translating measurements performed on monocultures to derive accurate models of a mixed co-culture’s behaviour, methodology for dynamic estimation of composition for a (non-engineered) co-culture based on multiple in situ population-averaged measurements of its behaviour (1B), and demonstration that when combined these approaches can be used to dynamically control a co-culture composition or stabilise its makeup for 7 days (*∼*250 generations), with the experiment ending before escape mutations overcome control.

## Results

We implement cybernetic control over the composition of a 20 mL *P. putida* and *E. coli* co-culture grown in a Chi.Bio [21], an automated bioreactor with heating, liquid handling, and fluorescence spectrometry capabilities. To achieve this control, we first characterised the bacteria’s fluorescence outputs under different conditions, used to inform composition estimates. We then characterise their growth rate and dynamics over a temperature range, showing that small adjustments can actuate changes in co-culture composition. Combining the two species into co-cultures grown at low or high temperatures can favour one species over the other as expected, but with undesirable behaviours that hinder robust control. After overcoming these challenges, we use the experimental data to build a system model and state estimator that together are used to demonstrate control.

We initially used the *P. putida* KT2440 strain in our experiments, but later swapped to instead use a *P. putida* OUS82-derived strain to improve estimation and control. This strain has a *lapA* gene knockout which reduces its ability to attach to surfaces and form biofilms [22, 23]. All results presented in this work (i.e. characterisation, control experiments) that mention *P. putida* refer to and are derived from OUS82, except for those in the ’Challenges in Co-Culture Implementation’ section which specifically highlight the issues with KT2440.

### Measuring Co-culture Composition

We began by identifying properties of our *P. putida* and *E. coli* co-culture that could be exploited to estimate its composition. With a view towards bioprocess applications where integrating flow cytometers or microscopes with bioreactors may be costly or impractical, we instead looked towards bulk culture OD and fluorescence measurements, which bioreactors can measure *in situ*. The Chi.Bio bioreactor [21] measures OD at 650 nm every minute, and maintains the co-culture at an OD setpoint (i.e. functioning as a turbidostat) by routinely diluting the culture with fresh media when it rises above a setpoint. The OD also contains information about the co-culture composition in how it changes over time (total growth rate - see State Estimation section below).

To acquire more online information on the composition, we identified differential fluorescence behaviours of both strains (as stationary phase monocultures) via a 2D fluorescence scan in a plate reader (Fig 2A). This highlighted an emission peak for *P. putida* at *∼*460 nm when excited at *∼*410 nm which is characteristic of the fluorescence of pyoverdine, an iron-chelating siderophore produced by the *Psuedomonas* genus [24]. *E. coli* had negligible observable fluorescence in this region, though some response is observed at excitation/emission wavelengths of 420 nm/500 nm. As the bioreactors can measure this fluorescence signal by exciting the co-culture at 395 nm and measuring emissions at 440 nm (the closest wavelengths available in the Chi.Bio), we showed that *P. putida* was producing a fluorescent molecule (assumed to be pyoverdine) at a concentration sufficient to be observed, and from its absolute value and dynamics could derive information on the amount of *P. putida* in the co-culture.

**Fig 1.**
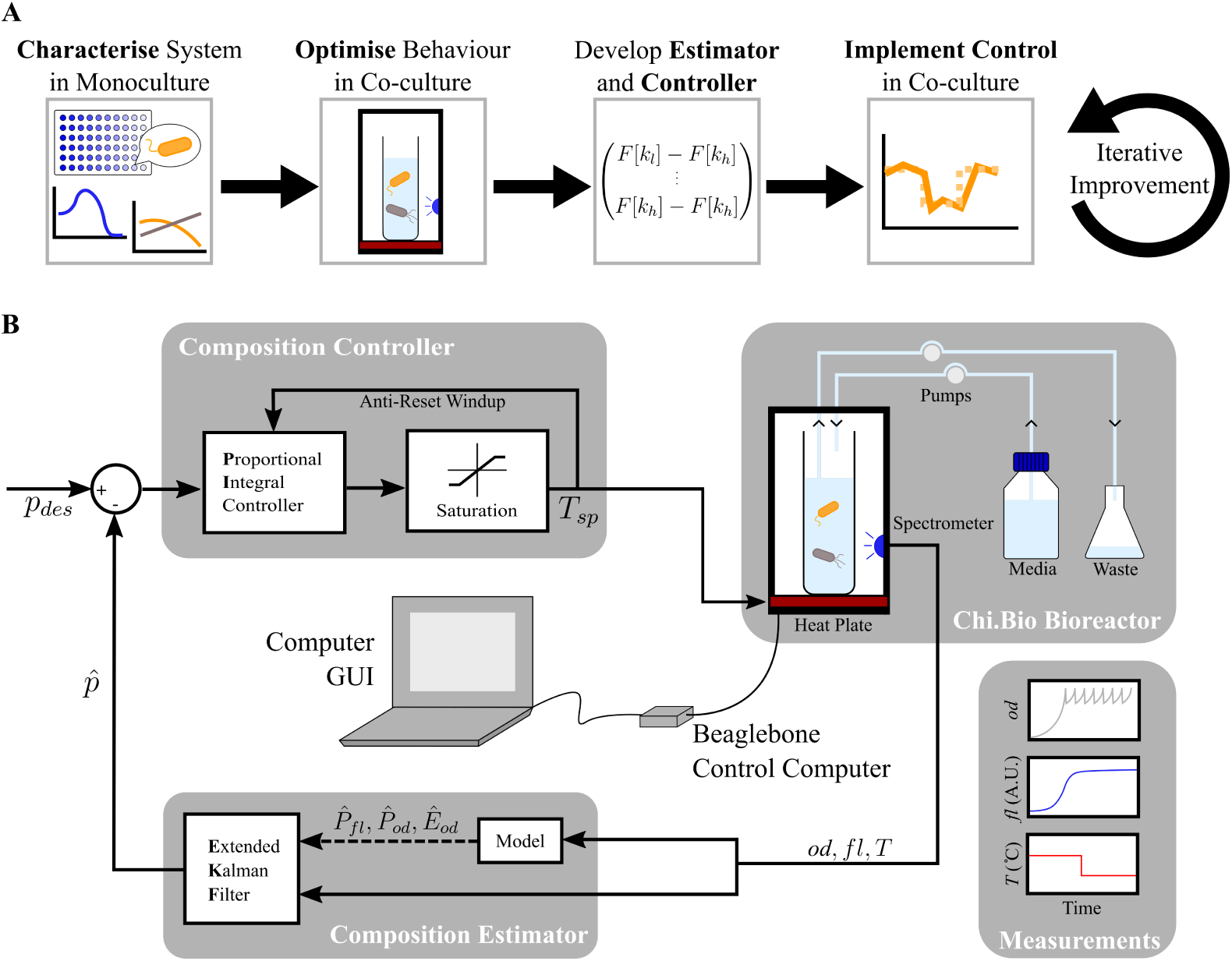
Cybernetic control approach and implementation. **(A)** Process of implementing cybernetic control of a co-culture. Begin by identifying suitable inputs and outputs to the system in monocultures, then combine both species and optimise the co-culture setup for better observability and controllability of the composition. Derive and parameterise a composition estimator and controller, then integrate them into the experimental setup. Finally, iteratively improve the setup, model, and control in co-culture experiments. **(B)** Automated cybernetic control of a *P. putida* and *E. coli* co-culture. The bacteria are cultured in Chi.Bio bioreactors functioning as turbidostats, where pumps dilute with fresh media and pump out waste. Spectrometers and temperature sensors take online measurements of bulk culture optical density (OD) *od*, fluorescence *fl* and media temperature *T* every minute, then feed these measurements into the control computer where a model of the system and an extended Kalman filter (EKF) derives an estimate of relative *P. putida* abundance *p*^. The difference between *p*^ and desired *P. putida* abundance *p_des_* is used by a proportional-integral (PI) controller to calculate a culture temperature setpoint *T_sp_*that would drive the co-culture towards the desired composition. A saturation block prevents unsuitable temperature setpoints and an anti-reset windup decreases integral buildup. The bioreactors’ heatplates alter culture temperature to the new *T_sp_*, affecting *P. putida* and *E. coli* growth rates and thus composition, and the process is repeated the next minute.

**Fig 2.**
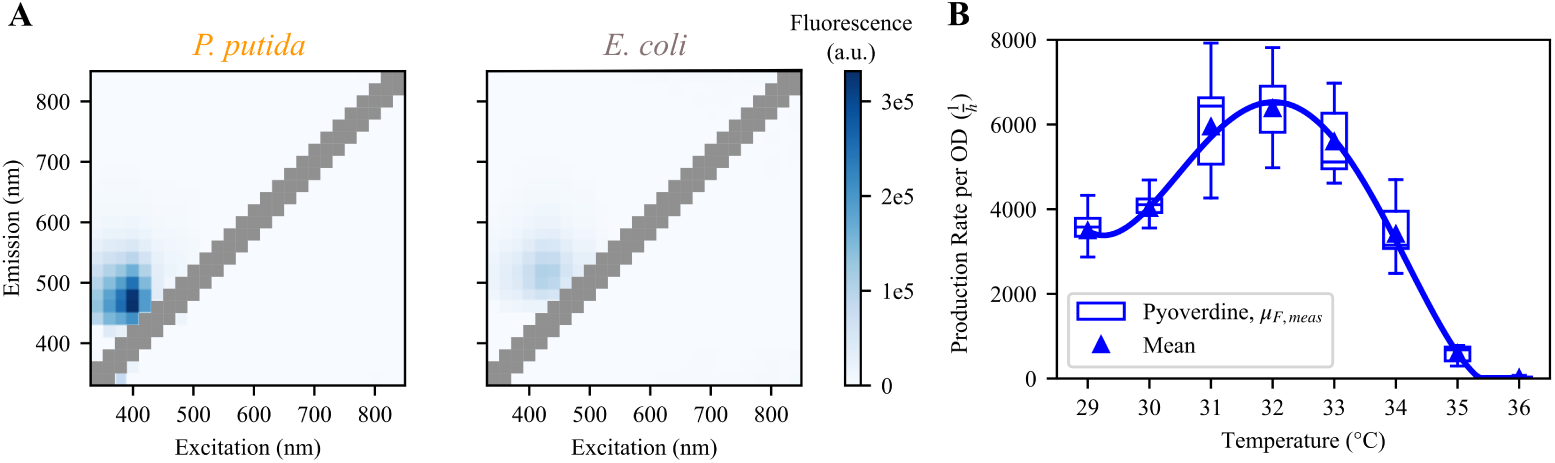
Fluorescence characteristics. **(A)** 2D excitation-emission fluorescence scan of *P. putida* (left) and *E. coli* (right) grown to stationary phase in a 96 well plate. *P. putida* naturally produces pyoverdine, a fluorescent siderophore with peak emission around460 nm when excited around 410 nm. At those wavelengths *E. coli* has an observable but insignificant amount of fluorescence. **(B)** Production rate of pyoverdine (h^-1^) at different temperatures peaking at 32 *^◦^*C, calculated by measuring fluorescence of a *P. putida* monoculture. The fourth-order polynomial fit is calculated using the mean value at each temperature.

When compared to engineered fluorescent reporters (which can be expressed by constitutive promoters), a weakness of naturally produced fluorescent molecules is that they may exhibit complex behaviours that depend on the host cell’s state. Using such a reporter as a measurement tool therefore requires characterisation of its dynamics under the range of conditions anticipated during experiments - which in this work meant understanding how pyoverdine production varied as a function of temperature. We did this by estimating pyoverdine production from the fluorescence of *P. putida* monocultures grown at different temperatures, accounting for how different growth rates affected the dilution of pyoverdine molecules, and observed that production peaks near 32 *^◦^*C and decreases at temperature extremes. Knowing this temperature-pyoverdine relationship, we have an additional parameter that can be used to infer *P. putida* abundance from current fluorescence values and historic temperature setpoints.

To ensure that fluorescence readings were comparable between reactors the calibration parameters for each reactor were measured (representing a reactor’s characteristic fluorescence offset and scaling factor respectively, S1 Figure), which also confirmed that fluorescence scaled linearly with the concentration of pyoverdine within the range relevant to our experiments. We also showed that changing culture temperature does not affect the fluorescence of pyoverdine in media with no cells, supporting the hypothesis that changes in observed fluorescence are due to increased pyoverdine production rather than any temperature dependence of measured fluorescence from a fixed quantity of pyoverdine S2 Figure.

Finally, to independently measure a “ground truth” co-culture composition during control experiments, an alternative offline measurement was required. This was initially attempted by routinely sampling from the co-culture and plating diluted samples onto chloramphenicol and antibiotic-free plates, as *P. putida* is naturally resistant to chloramphenicol (S3 Figure) (where *E. coli* was estimated as the total number of colonies subtracted by the number of *P. putida* colonies). However, while this method could show changes in orders of magnitude, even after some optimisation it remained too variable to allow discrimination of small composition changes (S4 Figure). To address this we employed an *E. coli* strain that contains a red fluorescent protein (RFP) cassette, used only for validating control in flow cytometry after bioreactor runs have already concluded. This was necessary because samples were stored at *−*80 *^◦^*C, and we observed that the fluorescence signal was lost after a freeze-thaw (S5 Figure), which prevented consistent flow cytometric gating and was hypothesised to be due to pyoverdine leaking out of cells. The RFP reporter is not used at any point for *control* of the co-culture, and is too weakly expressed (PRNA1 promoter) for any signal to be observed in a Chi.Bio when excited at its peak wavelength.

### Actuating Change in Co-culture Composition

Once composition could be measured, we needed a way to adjust co-culture composition to direct it towards desired reference compositions. Possible methods include tuning the concentration of chloramphenicol (increase to favour *P. putida* and vice versa) or a carbon source which only one species can utilise (e.g. lactose, only metabolised by *E. coli*), but for ease of implementation and broad applicability, we chose to exploit their different temperature niches. Previous work [25] has already shown that temperature can be used to influence the final composition of a batch culture of these species as *P. putida* grows optimally at a lower temperature than *E. coli* .

To characterise the relationship between temperature and growth rate under our experimental conditions (Fig 3A), *E. coli* and *P. putida* were grown in monocultures at a set temperature for 12 hours. The reactors act as a turbidostat (continuously diluting the culture) that ”dithers” the OD from 0.44-0.56 (S7 Figure), i.e repeatedly allowing the OD to increase until 0.56 before diluting it down to 0.44 with fresh media.

**Fig 3.**
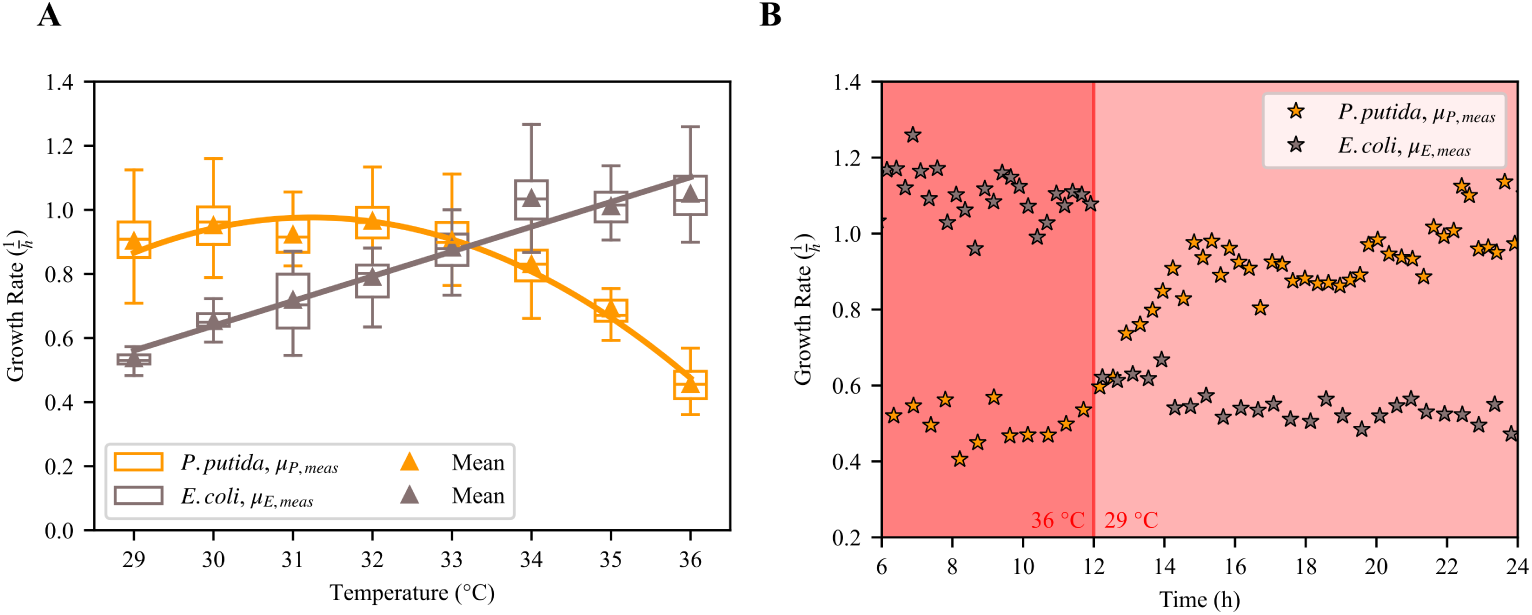
Composition actuation. **(A)** Growth rate of *P. putida* and *E. coli* (h^-1^) at temperatures between 29 to 36 *^◦^*C in M9 media supplemented with 0.02 % casamino acid (CAA) when held at a turbidostat OD setpoint of 0.5. A second-order (*P. putida*) and first-order (*E. coli*) polynomial fit is calculated using the mean value at each temperature. **(B)** Growth rate dynamics of *P. putida* and *E. coli* calculated from monocultures, where dark red background = 36 *^◦^*C, light red background = 29 *^◦^*C. While the *E. coli* growth immediately responds to the temperature change by dropping from 1.1 h*^−^*^1^ to 0.6 h*^−^*^1^, *P. putida*’s growth rate responds gradually, only reaching 1.0 h*^−^*^1^ after about 2.5 hours.

Afterwards, the temperature is swapped to a different setpoint for 12 hours to observe the growth response to a change in temperature. *P. putida* grows quickest at around 31 *^◦^*C and slower at either temperature extreme while *E. coli* ’s growth rate increases linearly with temperature. At around 33.2 *^◦^*C their growth rates intersect, yielding a temperature setpoint where *theoretically* a co-culture of any composition could be maintained. In contrast, temperature setpoints above or below this critical point favour the growth of *E. coli* or *P. putida* respectively. Temperature increases are rapid with the Chi.Bio’s heatplate actively heating the culture while temperature drops are slower, relying only on passive cooling (timescales dependent on the magnitude of change).

While investigating the time-dependence of our growth-temperature relationship we observed the two species respond with different time scales when the temperature changes: the *E. coli* growth rate drops sharply when the temperature changes from 29 to 36 *^◦^*C (Fig 3B), while the *P. putida* growth rate gradually decreases for around two and a half hours before settling. The same dynamics are observed (S8 Figure) when the temperature transition is reversed (i.e. cold to hot), indicating that this lag is specific to the species and not the direction of temperature change. Finally, our characterisation approach implicitly assumes the temperature-growth relationship in monoculture will be similar to that in co-culture - this was not obvious *a priori* as factors such as density dependence or inter-species interactions could lead to different dynamics in co-culture. Nevertheless, subsequent sections show this assumption *was* adequate for our application, though this is aided by the use of closed-loop feedback which reduces sensitivity to system parameters.

### Challenges in Co-culture Implementation

After initial characterisation in monoculture, we investigated the effect of temperature on a co-culture. This was initially done with the *P. putida* KT2440 strain. Monocultures were mixed and grown at either 26 *^◦^*C or 37 *^◦^*C (Fig 4A) - as expected 26 *^◦^*C strongly favoured *P. putida* KT2440 with no *E. coli* observed after less than 4 hours. However, this experiment demonstrated that despite *E. coli* ’s significant growth rate advantage at 37 *^◦^*C, *P. putida* KT2440 persists in the co-culture, and after an initial fall in abundance even seems to gradually increase over time.

**Fig 4.**
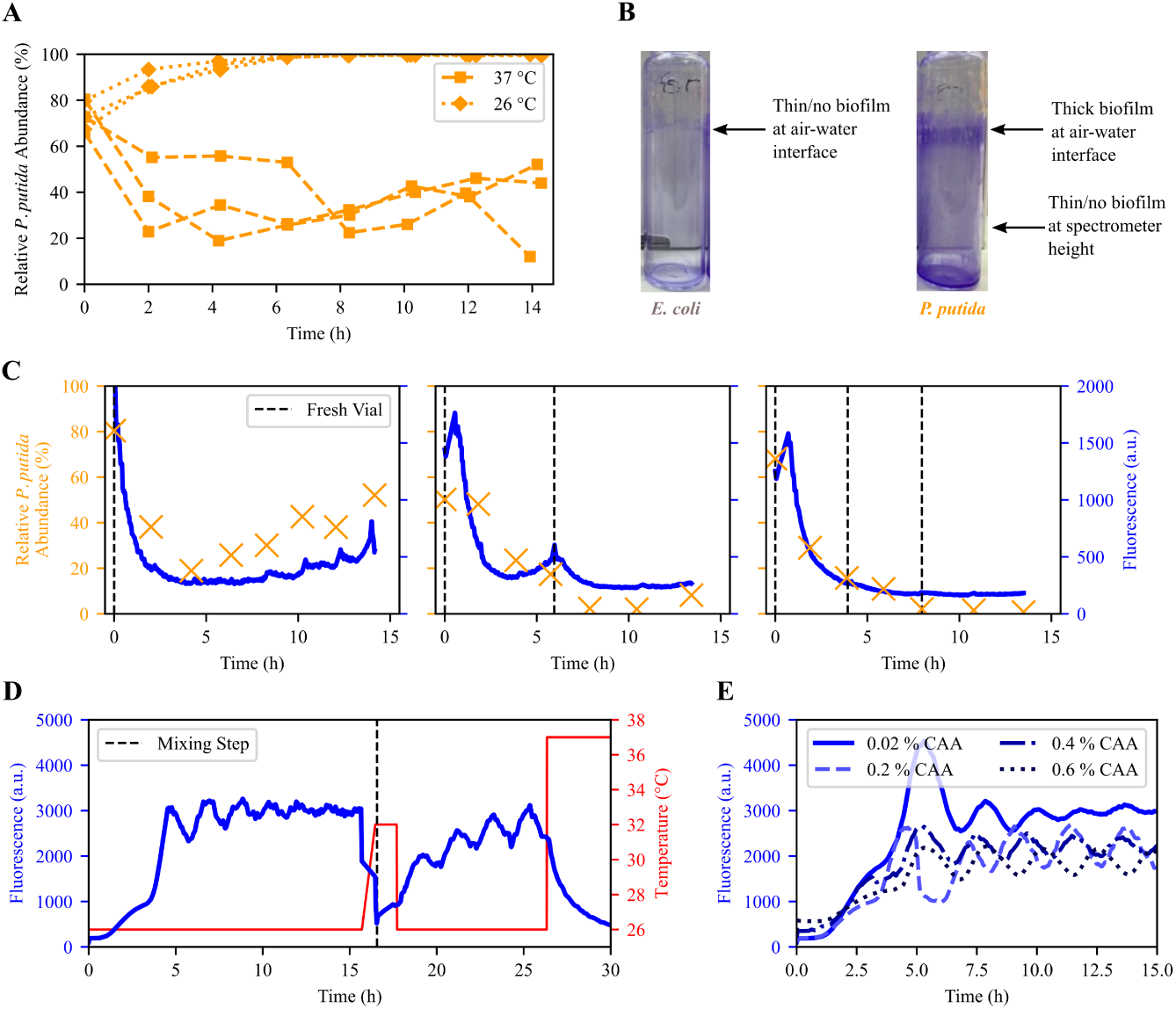
Challenges in Co-culture Implementation. **(A)** Relative abundance (%) of *P. putida* KT2440 and *E. coli* after mixing at *t* = 0 into a co-culture. At 26 *^◦^*C, *P. putida* KT2440 quickly and completely takes over the culture, while at 37 *^◦^*C *E. coli* is favoured but *P. putida* KT2440 persists even after 14 hours. **(B)** Examples of biofilms (stained with 0.1 % crystal violet) in vials from *E. coli* (left) and *P. putida* KT2440 (right) overnight monocultures. The dark purple near the air-water interface of the *P. putida* KT2440 vial indicates thick biofilm, while there is little/no biofilm at spectrometer height for both. **(C)** Relative *P. putida* KT2440 abundance (%) and fluorescence of co-cultures mixed at *t* = 0 and held at 37 *^◦^*C. (Left) Without changing vials, *P. putida* KT2440 biofilms slowly accumulate, artificially inflating *P. putida* growth and abundance, which in turn increases fluorescence. (Centre) Replacing the vial when fluorescence increased after about 6 hours was sufficient to remove most *P. putida* KT2440 from the co-culture. The gradual rather than sharp decrease in fluorescence in the fresh vial indicates that the rising fluorescence was from a genuine increase in pyoverdine rather than biofilms obscuring the spectrometer. (Right) By replacing vials every 4 hours (i.e. before biofilms can form), the *P. putida* KT2440 is completely lost from the co-culture, as expected by its lower growth rate at 37 *^◦^*C. **(D)** Fluorescence (a.u.) and temperature (*^◦^*C) of a *P. putida* OUS82 monoculture, mixed 50:50 with an *E. coli* monoculture at *∼*16 hours. At 26 *^◦^*C, the monoculture fluorescence first increases to around 3000 a.u. as the culture reaches the OD setpoint (not shown), and then oscillates while slowly dampening over time. Upon mixing, fluorescence drops sharply by half (as half of the *P. putida* is replaced with *E. coli*) and continues to decrease as the temperature is set to an intermediate temperature of 32 *^◦^*C. When the temperature is reduced to 26 *^◦^*C, fluorescence rises back towards 3000 a.u. while oscillating. **(E)** Fluorescence (a.u.) of *P. putida* OUS82 monocultures over time when grown in M9 with a range of CAA concentrations at 26 *^◦^*C. Oscillations are present at all concentrations, but while 0.02 % has a large spike upon reaching the OD setpoint, oscillations dampen over time, while at all other concentrations, they persist at the same amplitude over 15 hours.

Upon further investigation, we observed a biofilm concentrated around the air-water interface that grew in co-cultures and *P. putida* KT2440 monocultures (especially visible when stained with 0.1 % crystal violet, Fig 4B), but not *E. coli* monocultures. This suggests that the co-culture biofilm was primarily comprised of *P. putida* KT2440. The biofilms are not particularly thick around the bottom third of the tube where the OD and fluorescence measurements are taken. Additionally, transferring the culture to a fresh vial without biofilms did not significantly impact measurements (Fig 4C middle). Hence, we assumed that the biofilm was not significantly influencing the co-culture *measurement* through aberrant readings. Instead, we hypothesised that forming a biofilm granted *P. putida* KT2440 a strong selective advantage in the turbidostat environment as they remain in the bioreactor while planktonic cells are constantly pumped out as the culture is diluted. These biofilm cells would be able to divide and then shed back into solution, artificially increasing the overall *P. putida* KT2440 growth rate and allowing it to persist at temperatures where it should be quickly outcompeted by *E. coli* . We tested this by comparing a co-culture grown at 37 *^◦^*C with no vial changes (Fig 4C left), with a co-culture swapped into a fresh vial when fluorescence began to increase (i.e. when a biofilm had been established, Fig 4C middle), and with a co-culture swapped into a fresh vial every 4 hours (i.e. before a biofilm had been observed to form, Fig 4C right). We observed that after the initial post-mix drop in fluorescence and *P. putida* KT2440 abundance, both fluorescence and abundance began to increase in the reactor with no fresh vial. Meanwhile, swapping into a fresh vial when fluorescence began to increase could reverse the trend, and swapping every four hours prevented either fluorescence or abundance from recovering.

Attempting to mitigate short-term biofilm formation, we initially tested several anti-biofilm measures reported in the literature including Polysorbate 20 (Tween) [26] and cellulase [27]. While some were effective in standard biofilm assays in 96 well plates, they tended to be ineffective for cultures in Chi.Bio reactors, highlighting how different methods of culturing can create different phenotypes (S9 Figure, S10 Figure). This could be due to the overall cell density (cells reach a dense stationary phase in 96 well plates, while they are held in an early exponential phase in the Chi.Bio) or the fact that constant dilution in turbidostats selects for cells in biofilm regardless of chemical/biological intervention. We therefore changed from *P. putida* KT2440 to instead use *P. putida* OUS82 with a *lapA* gene knockout (the characterisation data in the above section is for this strain, and it is used for all subsequent co-culture experiments). *lapA* encodes a surface adhesin described as “essential” for biofilm formation [22, 23], but even so, biofilms still appear after *∼*24 hours in *P. putida* OUS82 monocultures (or longer in co-cultures, depending on the population ratio). Because of this, cultures were transferred into sterilised, fresh tubes every 24 hours.

While the above mitigations minimised the impact of biofilms on co-culture dynamics, parameterisation of OUS82 revealed unexpected fluorescence dynamics unseen in KT2440: robust oscillations in pyoverdine production emerged that were highly consistent across replicates and persisted for days even after mixing in co-culture (Fig 3D). These oscillations have a significant impact on the ability to estimate *P. putida* abundance and thus on control, as an increase in *P. putida* population is indistinguishable from the rising edge of an oscillation. Further experiments (S11 Figure) showed these oscillations were affected by temperature, becoming more prevalent at lower temperatures. This makes accurate calculation of the mean pyoverdine production rate challenging unless the entirety of an oscillation period had a constant temperature. The oscillations were also influenced by the concentration of CAA (Fig 3F), particularly arginine, which is known to be related to pyoverdine production [28]. The effect of several synthetic amino acid mixtures, nitrogen sources, and arginine concentrations either in media or spiked-in were tested (S12 Figure, but could not completely remove oscillations at 26 *^◦^*C.

Ultimately, the simplest set of conditions that produced consistent results was 0.02 % CAA and a minimum temperature of 29 *^◦^*C. Reducing CAA concentration dampened oscillations quicker (Fig 3F) at all temperatures, and staying above the minimum temperature prevented excessive oscillations. Static references were also less affected, as they depend on reaching a particular composition and then maintaining it with a 33.2 *^◦^*C critical point where little-to-no oscillations are observed. Removing CAA entirely was not practical because *E. coli* grows too slowly in comparison to *P. putida* without it (S15 Figure). The limited temperature range of 29 to36 *^◦^*C was therefore chosen for actuation of the co-culture to trade off different behaviours of the combined system: at lower temperatures the larger difference in growth rate (and quicker control response) would be offset by the larger oscillations, making composition estimation difficult. On the other hand, higher temperatures inhibit pyoverdine production (Fig 3A), again reducing our ability to accurately track composition.

### State Estimation and Model

The complex dynamics of pyoverdine production (i.e. temperature dependence, oscillations) make bulk culture measurements of absolute fluorescence by itself an inaccurate direct proxy for *P. putida* abundance. A simple example is that a co-culture that is 99 % *P. putida* but held at a high temperature for several hours will have almost no measurable fluorescence, despite consisting primarily of *P. putida*. To overcome this challenge, we combine several estimations derived from easily available bioreactor measurements to achieve an accurate estimation of composition:

1. *Absolute OD value*: the combined abundance of both species
2. *Time derivative of OD*: co-culture growth rate over time
3. *Absolute fluorescence value*: the abundance of pyoverdine
4. *Time derivative of fluorescence*: the production of pyoverdine over time

The absolute OD value (1) reflects the total abundance of cells. This requires calibration (e.g. against CFU or flow cytometry) if a measure of the *number* of cells for each species is required (due to differences in size and absorption between species leading to different numerical densities for a given optical density). The cultures are grown in turbidostat mode (i.e. maintained around a set density), and so are periodically diluted when the absolute OD exceeds its setpoint, creating phases of dilution and phases of growth (S7 Figure) - we refer to each such period between dilution events as a *growth period*. The time derivative of OD (2) is a composite measure of the two species’ growth rates over the last growth period. Since the temperature at every time point is known, as is the individual species’ growth-temperature relationship, the time derivative of OD can provide the abundance estimates, *P*^^^*_od_*[*k*] and *E*^^^*_od_*[*k*] (S1 File). As an example, a co-culture grown at 36 *^◦^*C with a high total co-culture growth rate (i.e. equal or similar to *E. coli* growth at that temperature) indicates a predominance of *E. coli* , while one with a low total co-culture growth rate should consist predominantly of *P. putida*. The absolute fluorescence values (3) provide the next source of information about composition through the relationship shown in Fig 2B, though as described above this provides limited information in certain conditions. Finally, the time derivative of fluorescence (4) is the function of historical time-averaged *P. putida* abundance and the temperature, and given an understanding of the temperature-production relationship, can be used to infer the abundance estimate *P*^^^*_f l_*[*k*] (S1 File).

Estimation method 1 reflects *total* density of the culture while estimation methods 2-4 provide an estimate of composition, each with distinct advantages and disadvantages (Table 1). To reduce the effect of measurement noise, methods based on time derivatives require multiple consecutive measurements. This results in a lower measurement update rate and reduces their accuracy during periods in which temperature changes rapidly. More broadly, the quality of all estimation methods depends strongly on the magnitude and dynamics of culture temperature. For example, the quality of estimates from absolute fluorescence decreases at low temperatures when oscillations begin to appear, as well as at high temperatures where pyoverdine production is minimal (i.e. above 35 *^◦^*C, see Fig 3A). Curvature methods (particularly OD, which does not oscillate) remain effective at low and high temperatures, however, the accuracy of OD curvature measurement methods decreases whenever a culture is around the 33.2 *^◦^*C critical temperature, as (near) equivalent *P. putida* and *E. coli* growth rates mean the total co-culture growth rate becomes independent of composition. Together these comparisons highlight the complementary strengths and weaknesses of each estimation method, which when taken together demonstrate that at least one method is viable in each condition encountered by our co-culture.

**Table 1.**
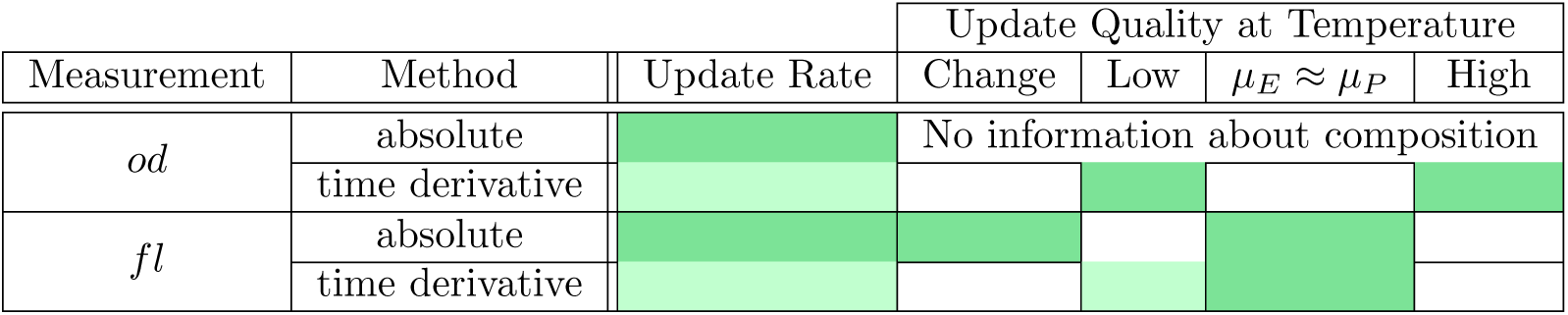
Qualitative comparison of estimation methods or co-culture composition. . Dark green represents a high information update rate/quality, light green intermediate, and white low or no useful information.

In order to combine each method to derive a single quantitative estimate of composition, which also accounts for each method’s variable quality (and its dependence on the culture’s current and past behaviour), we employ an EKF, a well-established control engineering tool that estimates the state of a system by fusing a system model with measurements and their uncertainty. Several works have explored how it can be used to improve observability of biological systems [29–31], here we demonstrate it experimentally: every cycle (i.e. every minute), the abundance of *P. putida P* [*k*], *E. coli E*[*k*], and pyoverdine *F* [*k*] are calculated in two steps: the prediction step and the measurement update step (Fig 5).

**Fig 5.**
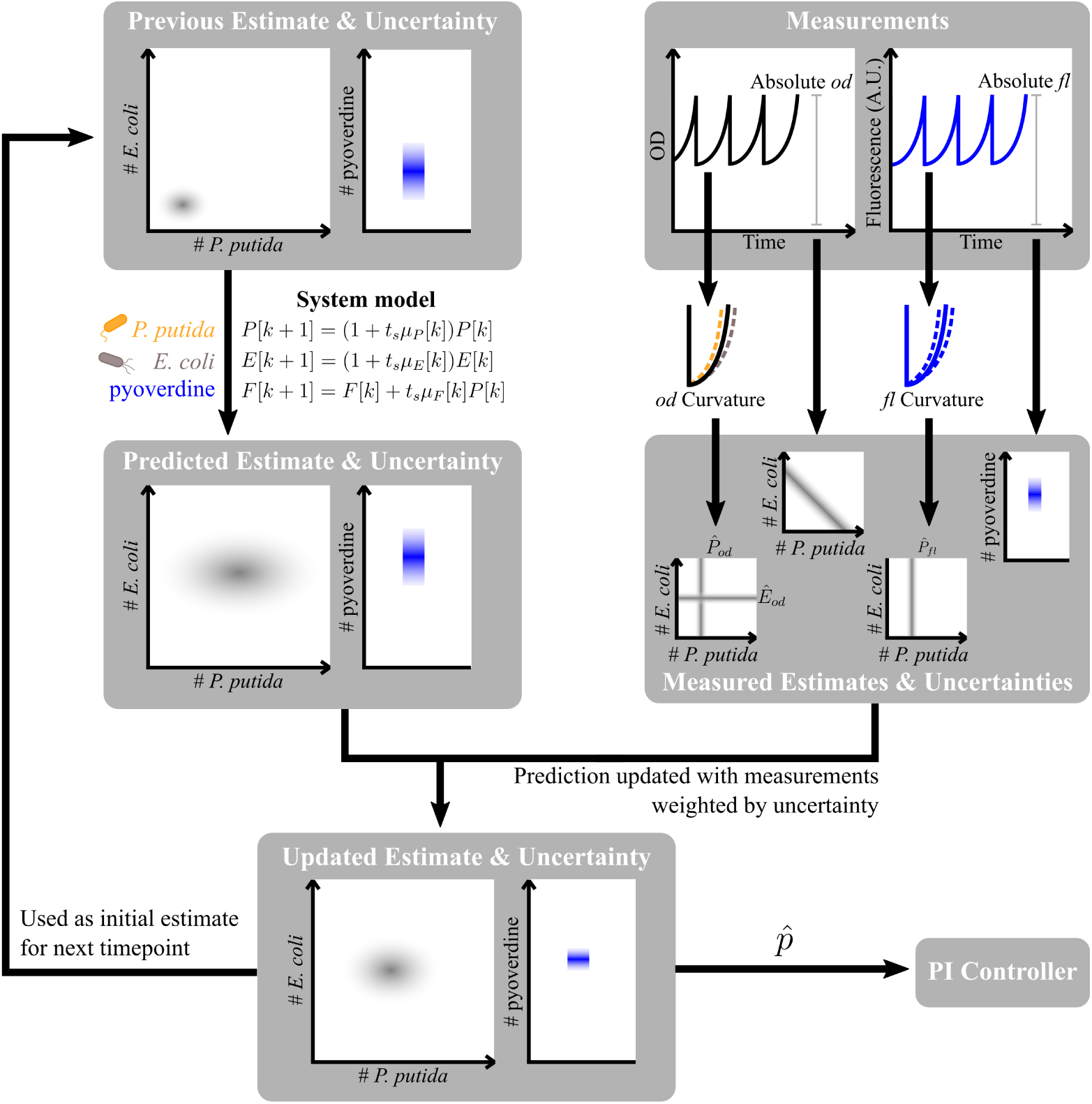
Extended Kalman filter for recursive online composition estimation. First, the current abundances of *P. putida*, *E. coli* , and pyoverdine are predicted based on the estimate from the previous cycle and the system model. In the next step, the bioreactor takes bulk culture fluorescence and OD measurements, whose curvature across a growth period is used to calculate the intermediate estimates *P*^^^*_od_*, *E*^^^*_od_* and *P*^^^*_f l_*. The EKF then updates the predicted abundances with the intermediate estimates and the absolute measurement values, taking into account the model and measurement uncertainties. Finally, from the resulting estimate, the relative abundance of *P. putida p*^ is computed and used by the controller.

In the prediction step, the EKF uses an initial state estimate from the previous cycle, combining this with a system model to predict the current state of the system. The system model is described in detail in S1 File; it uses differential equations to model the dynamics of each species’ growth and pyoverdine production and is kept sufficiently simple (i.e. with a small number of variables and parameters) to facilitate system characterisation. While this model *could* be used to make long-term forward predictions of the system state based only on an initial state estimate, any errors in the initial estimate or the model itself would be propagated forward, leading to divergence of the model from reality. Overcoming this divergence is the purpose of the measurement update step; here the EKF updates the predicted state with the different measurements and intermediate estimates, weighted by their uncertainty so that low-quality updates (e.g. *P*^^^*_f l_*[*k*] at a high temperature) contribute less to the final estimate. This allows the individual measurement update methods to complement each other and makes the resulting updated state estimate less prone to measurement and system model inaccuracies.

The capabilities and limitations of this approach are explored in Fig 6. A realistic simulation including plausible measurement noise and model mismatch (Fig 6A) demonstrates that while OD based estimates *P*^^^*_od_*[*k*] and *E*^^^*_od_*[*k*] are helpful in the beginning, they were far away from the true composition toward the end of experiment when culture temperature was near to the critical temperature. Meanwhile, using only fluorescence in the measurement update step was insufficient when the temperature was at 36 *^◦^*C near the beginning, but delivered good results afterwards. Finally, we observe that the state estimate without any measurement update has no opportunity to correct for its model mismatch and (simulated) noise, and continues to diverge over time. The result of this simulation is borne out in summary statistics from experimental co-culture control tests (Fig 6B), where the EKF with combined updates leads to more accurate composition estimates than the system model alone. This is observed in both short-term and long-term experiments, with the latter presenting a broader error distribution due to increased time for estimation error to build up. It is worth noting that due to biological and measurement variability during experiments, one anticipates cases in which the model-only approach is (by chance) closer to the actual density than the EKF output; nevertheless, the EKF approach outperforms model-only prediction at almost all time points in almost all experiments.

**Fig 6.**
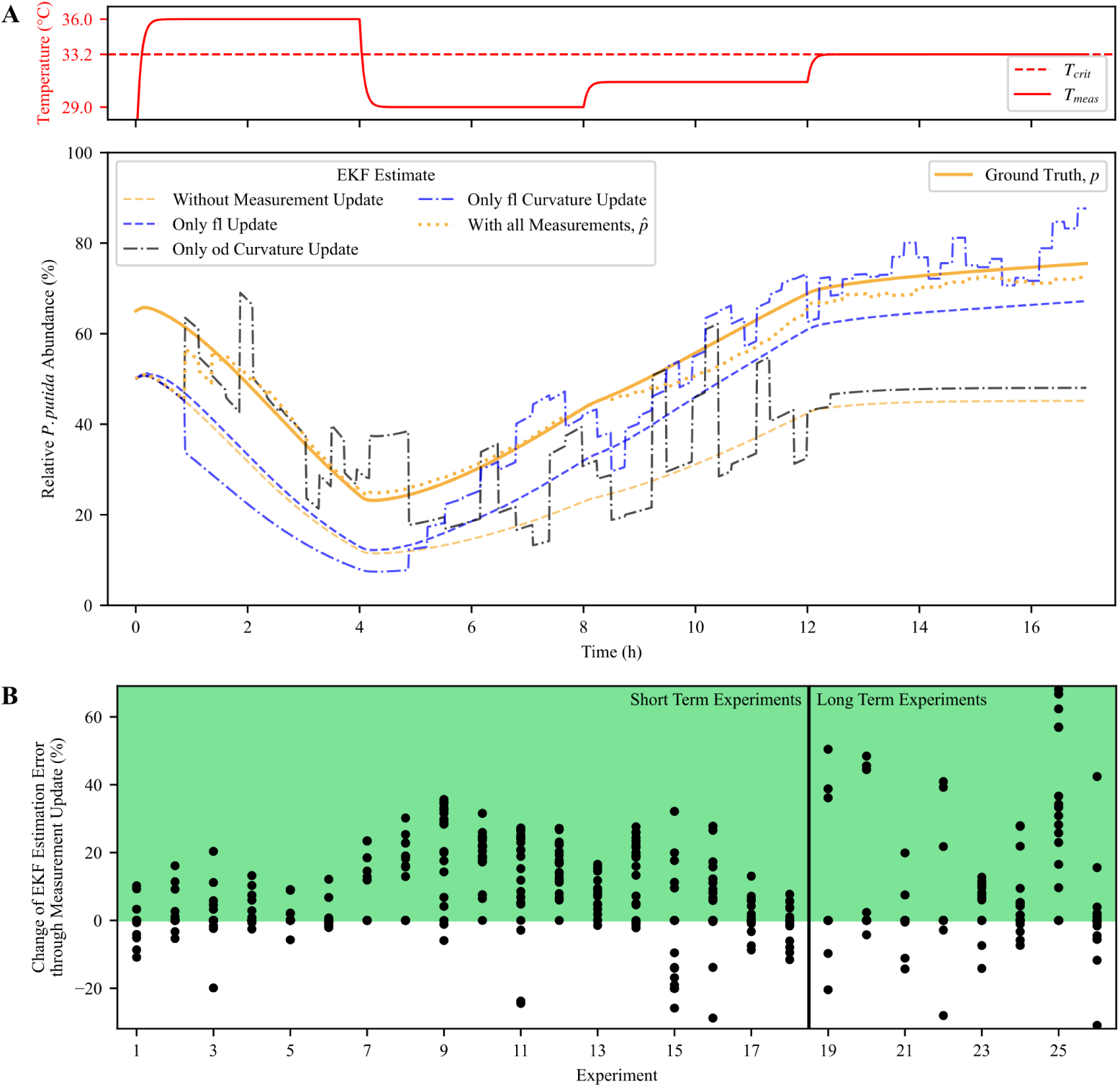
Comparison of different extended Kalman filter composition estimates. **(A)** Composition of a co-culture including plausible measurement noise and model mismatch, simulated using the characterised system model (labelled as ground truth, *p*). (Top) The predefined temperature profile over 17 hours. (Bottom) The relative *P. putida* abundance estimates resulting from EKFs that employed either no measurement update, only a single update method, or all update methods combined. **(B)** Comparison between the error of the relative *P. putida* estimate from the EKF *p*^ and the error of the estimate from the system model alone (i.e. without any measurement updates) with respect to the offline flow cytometry measurements. The data was collected across 26 real co-culture experiments. A value above zero (green) signifies the EKF with all measurement update methods outperforming (i.e. reducing error) compared to the model alone. Long-term experiments span multiple days, while short-term experiments ran for *∼*1 day.

### Control Approach and Implementation

The system’s controller is tasked with setting the temperature such that the estimated relative *P. putida* abundance *p*^ shifts towards the desired relative abundance *p_ref_* . The family of proportional-integral-derivative (PID) controllers was chosen due to its computational simplicity and low model dependence [32]. The derivative term was not utilised as the PI controller demonstrated sufficient performance, and an over-damped (i.e. slow responding) control that introduces slower changes in temperature setpoints was found to improve state estimation. This fact also discouraged the use of a (computationally even simpler) Bang-bang controller which, given a dynamic system, usually results in zigzag-shaped temperature profiles. The output temperature was constrained to be within the 29 *^◦^*C to 36 *^◦^*C bounds. This constraint prevents large temperature fluctuations, preventing the controller from making swift adjustments that could otherwise lead to over/undershooting target community compositions, but comes at the cost of possible “integrator wind-up” in the control [32]. An anti-windup reset was therefore added that decreases the integrated error once the temperature setpoint saturates. The control gains were initially tuned in a simulation with final tuning being done in experiments. Both the EKF and controller were integrated into the Chi.Bio operating system in Python, running in real-time on the bioreactor’s control computer (a Beaglebone Black microcontroller).

By combining measurement, state estimation, a control algorithm, and actuation, we are able to implement cybernetic control of the *P. putida*-*E. coli* co-culture. All control experiments had an initial 1-hour period of growth at 33.2 *^◦^*C where the culture grew to reach the OD setpoint. We were able to demonstrate short-term static and dynamic reference tracking with a square wave pattern from 30 to 70 % *P. putida* abundance over one day (Fig 7A). The controller first drives the composition to the target and then maintains it by staying around the critical temperature with only slight deviations. The estimation improves over time and is able to align with offline measurements within 5 hours, negating the effect of variability in the initial inoculum mixing (i.e. when pre-cultures were combined to attain an approximately 50:50 population ratio), contrasting open-loop control which would propagate errors through the rest of the experiment. Temporary estimator inaccuracies (such as initial offset from 50:50 co-culture) do not significantly impair the control due to the relatively slow dynamics of the system. The sudden drop around 16 hours is assumed to be due to noise in the offline measurement process (storage/sample preparation/flow cytometry), as it is not consistent with adjacent time points given the temperature of the culture. We also demonstrate the response time of the controller with a sine wave reference (Fig 7B); here composition responds with a delayed and damped sine wave of a similar period. This is primarily due to the integrator in the PI controller and the slow system dynamics that result in a gradual and delayed change in the composition after a change in the reference.

**Fig 7.**
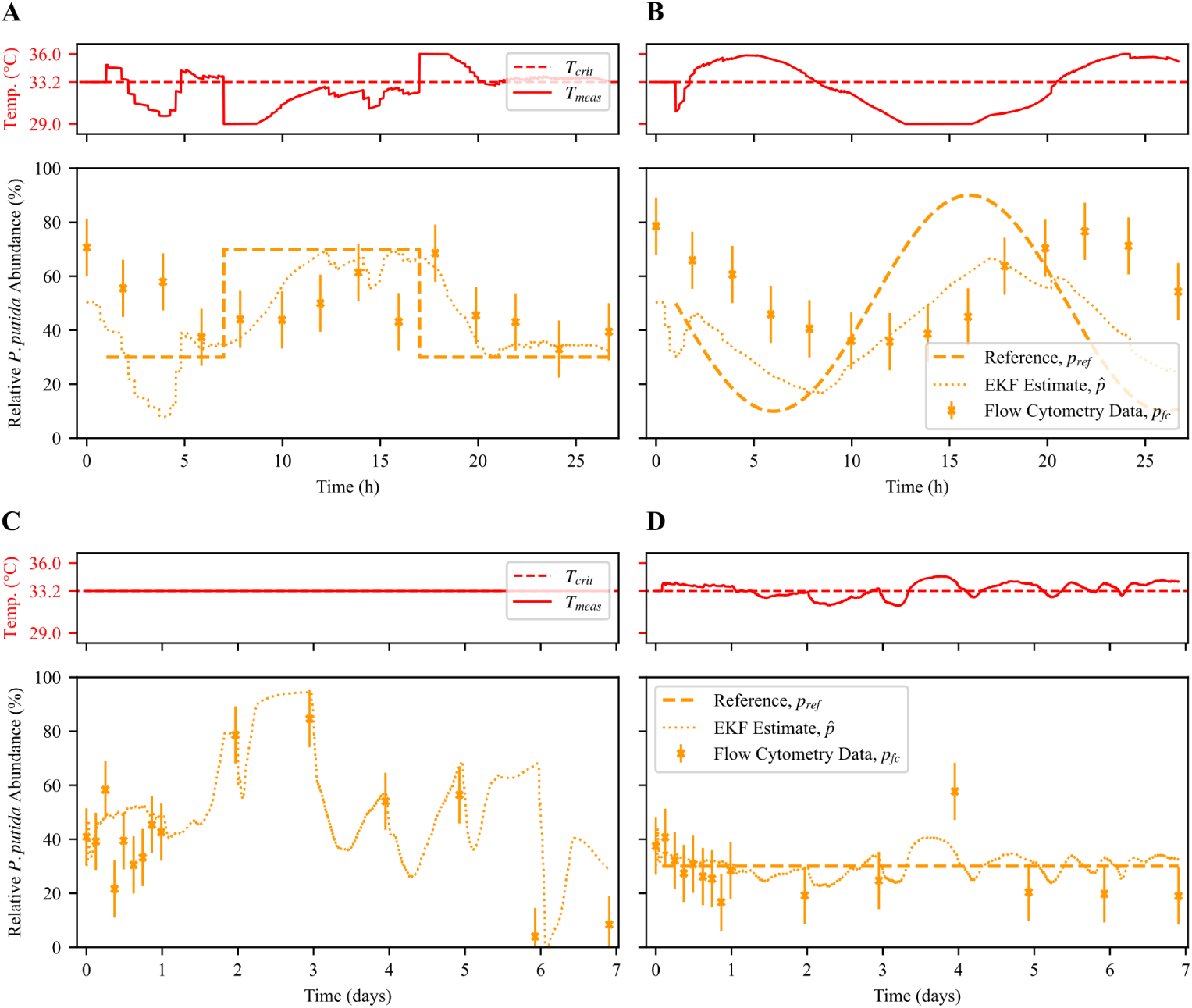
Open-loop and closed-loop control experiments. **(A)** (Top) Media temperature over time, as set by the controller. (Bottom) Relative *P. putida* abundance over time in a co-culture. Despite inaccuracy near the beginning (the estimator starts at an idealised 50:50 mix but mixing is imperfect) the EKF estimate improves over time and the controller is able to track a square wave reference. **(B)** Controller with a sine wave reference. As expected, the simple PI controller cannot fully compensate for the slow system dynamics and causes a delayed response. **(C)** Open-loop control of a co-culture over 1 week with starting ratio of 30:70 *P. putida*:*E. coli* grown at critical temperature 33.2 *^◦^*C, where theoretically the ratio would be maintained. Culture is swapped into fresh vials daily to prevent excessive biofilm formation, with samples for flow cytometry taken concurrently. While the composition is relatively static in the first 24 hours, it subsequently varies significantly (from 5-85 %). The EKF estimator tracks the composition well throughout the experiment. **(D)** Closed-loop control with the same 30:70 reference. The controller was able to maintain the *P. putida* abundance at around 30 % for the whole week, requiring slight deviations from the 33.2 *^◦^*C critical temperature in both directions.

While open-loop control of the co-culture *could* be possible given that the relationship between temperature and growth rate is known, without feedback such an approach would be unable to respond to phenotypic variation or adaptation, or indeed any genetic (evolutionary) adaptation that occurs during long-term growth experiments. For example, a co-culture mixed in a 30:70 ratio of *P. putida*:*E. coli* and held at the critical temperature should theoretically maintain that ratio as both species have an equivalent growth rate. This is tested in (Fig 7C), where the *P. putida* abundance fluctuated near 30 % for the first 24 hours, increased to *∼* 80 % after 2 days, and rose further before dropping to 10 % after 7 days. Fig 7D illustrates a similar experiment, but now employing the PI feedback control system with a set-point of 30 % for all 7 days. Excepting one outlier on day 4, the experiment shows that the set-point was maintained throughout, but with a consistent 10 % offset between the online estimated composition and the flow cytometric ground truth. This offset may be caused by the biofilms (of mixed composition) that formed in the vials even with daily vial replacement (S16 Figure): changes to the observed growth rate of either species (e.g. if it were to be artificially inflated by biofilm shedding cells into the media) lowers the quality of the OD curvature estimate. In addition to the possibility of systematic sampling noise in the flow cytometry protocol, flow cytometry only measures the composition of planktonic cells, completely ignoring biofilm cells which *do* contribute to EKF estimates as they both contribute to OD and fluorescence measurements.

Interestingly, while both biological replicates of long-term closed-loop control were able to maintain *P. putida* abundance at around 30 % (S17 Figure), they appeared to adapt differently and required the controller to compensate in opposite directions: one had an average temperature of 33.33 *^◦^*C (above the critical temperature), while the other had an average temperature of 32.35 *^◦^*C (below the critical temperature). This exceeds the noise of the temperature sensor, which has a base accuracy of *∼* 0.3 *^◦^*C. A range of phenotypic or genetic adaptations that affect measurement or actuation could be responsible: changes in the speed or dynamics of pyoverdine production, an increase in biofilm-forming ability, adaptation to a new growth temperature, adaptation to the media, adaptation to the other species, or more. In the timescales tested, the adaptation(s) have not imparted enough of a selective advantage to overwhelm the controller, which automatically compensates and maintains the composition at the desired reference.

## Discussion

This work demonstrated a cybernetic approach for controlling the composition of a natural two-species co-culture. While the interfacing of biological systems with computers to implement control has previously been described as cybergenetic [17, 19], it is often in the context of controlling the expression of genes [20]. To highlight that control of cellular behaviour is achieved without genetic programming, we describe this work as cybernetic (i.e. an automatic control system with feedback). By estimating composition from measurements of OD and natural fluorescence, computing control actions, and then changing environmental conditions to favour one species or the other, the setup can track static and dynamic reference compositions. In addition to advantages highlighted by other cybernetic works such as real-time reduction in variability, independence from inoculation/mixing ratios, the ability to set arbitrary references that can be changed mid-experiment, and the ability to compensate for adaptation over time, this approach attempts to extend cybernetic methods further, removing the need for genetically engineered parts to interface between cell and computer. It also does this with commonly available bulk-culture measurements instead of relying on automated microscopy, flow cytometry, or liquid handling/sampling equipment, which might be more costly and difficult to scale (in size of reactor or number of replicates).

Past cybernetic studies of co-culture control have often identified genetic mutation leading to loss-of-function as a key determinant of their longevity. In our system a similar challenge could emerge should the point of equal growth (i.e. intersection in Fig 3A) move beyond the range of allowable temperatures, or if the temperature-growth relationship for both species flattened significantly, reducing the coupling between changes in temperature and composition. This was not observed in any of our experiments, which maintained control for at least 7 days of exponential growth (*>*250 generations assuming an average species doubling time of 40 minutes), significantly longer than other cybernetically controlled bacterial co-cultures to-date. Nevertheless, past studies of microbial adaptation to temperature extremes indicate significant changes in growth optimum *are* possible [33, 34], though these would need to be very significant to eliminate our controller’s ability to adapt. More broadly, *all* exploited natural characteristics may drift over the course of long experiments, but combining several measurements, of which some are tied to the cellular growth rate (e.g. pyoverdine chelates iron, which is a crucial nutrient), and thus not having a single point of failure, shows the broader potential of EKF-type estimation approaches to provide robustness beyond what may be possible using a single fluorescent reporter or quorum sensing molecule. The reference ratio and its dynamics may also influence the accumulation of mutations-compared to directed evolution experiments which constantly and strongly select for a particular characteristic, a static reference should drive the composition to a particular ratio and then only maintain an environment that equalises the growth rates, an environment that should not impart as strong a selective pressure.

While our approach achieved reasonable control, several areas of improvement remain for future work. For example, growth rates were parameterised in monocultures at an OD of 0.5, where both bacteria are (approximately) in mid-exponential growth phase. However, in the co-culture, each species is at a lower OD (i.e. the sum of their ODs is 0.5), where the growth rate and dynamics might differ. In addition, as evidenced by the effect of temperature on pyoverdine production, control which alters a cell’s growth rate influences its physiological state, which in turn influences the cell’s bioproduction capabilities. Static references where the culture is held at the critical temperature would likely be less affected, as at that temperature *P. putida* and *E. coli* grow at 91 % and 81 % of their maximum growth rates, but a dynamic reference with temperatures closer to either end of the range might be more impactful. This tradeoff relies on the advantage of a robustly controlled co-culture outweighing the disadvantages of altered cell states.

Furthermore, we first attempted to use the *P. putida* KT2440 strain but because of its persistent biofilm formation (despite chemical and biological intervention), swapped to an OUS82 strain with a *lapA* knockout, which encodes a surface adhesin described as ”essential” for biofilm formation [22, 23]. While no reported conditions have previously been able to rescue *P. putida lapA*’s mutant biofilm defect [22], in our setup biofilms still appear after around 24 hours in *P. putida* monocultures or longer in co-cultures, depending on the population ratio. This is likely because turbidostat or chemostat setups provide a very strong selective pressure for biofilm-forming cells by constantly diluting out planktonic cells. While the controller can compensate for the effect of biofilms, it clearly affects the system during long-term experiments, favouring *P. putida* growth until swapping into a fresh vial, at which point only planktonic cells (predominantly *E. coli*) are transferred, resulting in a sharp drop in estimated and real composition (e.g. as observed in Fig 7C). Vial changes were done concurrently with sampling, providing an explanation for why estimated composition drops after most flow cytometry data points. However, while biofilm formation will continue to be an issue in larger, scaled-up bioreactors due to the potential for clogging or difficulty in sterilisation, its effect on control should be significantly dampened - the volume of culture scales by the cube, while surface area (and thus the relative amount of biofilm) scales by the square, decreasing the negative impact of biofilm on control with scale.

While demonstrated with *P. putida* and *E. coli* , this cybernetic control approach should be generalisable to other microbes by following a similar process of identifying methods of actuating and measuring culture composition, characterising them in monoculture, and then developing a model and controller. If using microbes with similar temperature niches, any other characteristics can be exploited to actuate composition including differing substrate utilisation [18], optimal nutrient niches (e.g. halophilic *V. natriegens* [35] actuated with salt concentration), natural resistance to chemicals and antibiotics (e.g. tetracycline resistant *B. subtilis* [36]), or sensitivity to light (e.g. cyanobacteria or microalgae). Similarly, while some other microbes also produce naturally fluorescent (e.g. chlorophyll in photosynthetic microbes) or luminescent (e.g. *A. fischeri* [37]) molecules that can act as reporters for composition measurement, employing an extended Kalman filter to combine with other measurements, the curvature and absolute value of OD is strain agnostic and can always be used.

Extending this approach to other species of interest should enable robust control of co-cultures for a wide range of biotechnological applications, effectively leveraging and unifying the most attractive properties of computational and biological systems.

## Materials and methods

### Strains and Media

Unless specified, the strains are *P. putida* OUS82 with a *lapA* gene knockout and *E. coli* MG1655 with a genomically encoded RFP cassette with a PRNA1 promoter. Some of the *P. putida* used in experiments shown in the ”Challenges in co-culture implementation” section was the *P. putida* KT2440 strain. Strains are stored in 25 % glycerol at *−*80 *^◦^*C and revived by streaking overnight onto LB-agar incubated at 30 *^◦^*C (for *P. putida*) or 37 *^◦^*C (for *E. coli*). Unless specified, experiments are done with M9 (Formedium) supplemented with 0.4 % glucose (Formedium), 2 mM Mg_2_SO_4_ (Sigma Aldrich), 0.1 mM CaCl_2_ (Sigma Aldrich), and trace elements (final concentrations of 50 mg L*^−^*^1^ EDTA.NA_2_.2H_2_O, 30.8 µM FeCl_3_, 6.16 µM ZnCl_2_, 0.76 µM CuCl_2_.2H_2_O, 0.42 µM CoCl_2_, 1.62 µM H_3_BO_3_, and 0.081 µM MnCl_2_.4H_2_O, all from Sigma Aldrich). For some experiments in the ”Challenges in co-culture implementation” it was supplemented with a range of CAA from Formedium, but unless specified all other experiments used a final concentration of 0.02 % CAA. For experiments exploring the effect of CAA or other amino acids on oscillations, arginine, serine, glutamine, and Complete Synthetic Mixture (CSM) with double dropouts for arginine and tryptophan (all from Formedium) were used. When spiking with amino acids, they are added to the same concentration in the reactor and media bottle.

### Fluorescence Scan

*P. putida* and *E. coli* were grown overnight in LB at 30/37 *^◦^*C, washed and resuspended in PBS (Formedium), and then 200 µL pipetted into the wells of a 96 well plate (Greiner Bio-One). This was measured in a Tecan Spark plate reader in ”Fluorescence Intensity 3D Scan Top Reading” mode with an excitation and emission wavelength range of 280 - 900 nm with a step size of 20 nm, bandwidth of 20 nm, manual gain of 100, and 30 flashes each.

### Bioreactor Setup

To benchmark 440 nm fluorescence measurements between reactors, each bioreactor was associated with a unique identifier and then calibrated with a dilution range of a filter-sterilised *P. putida* monoculture grown to stationary phase (removing cells which might scatter/absorb light, but retaining extracellular pyoverdine). The un-normalised fluorescence values were collected after stirring for 5 seconds and settling for 5 seconds, mimicking experimental conditions. For control experiments, the identifiers of the reactors being used were entered into the control computer, and the controller used the unique reactor offsets and scaling factors to calculate a normalised fluorescence value. For OD readings, before each experiment, all vials were autoclaved and wiped with ethanol, filled with media, heated to 33.2 *^◦^*C with stirring on, and then used as a blank.

To calculate growth rates for parameterisation or during control experiments, the OD of the culture was ’dithered’ around a setpoint of 0.5, where it was allowed to grow to 0.545 before being diluted to 0.455. The growth rate was calculated from the logarithmic slope between these two points. These values were selected instead of the default Chi.Bio dithering parameters to increase the frequency of dilutions, which improved the performance of the estimator. Parameterisation was done by growing for 12 hours at one temperature before swapping to another, and calculating the average growth rate after the culture fully adjusts to the new temperature.

3D-printed lids were used to cap duran bottles containing fresh media. To prevent negative pressure from pumping out media, they contain an ethanol sterilised tube with a 0.22 µm filter (Sartorious) that allows sterile air into the media bottles. The bioreactor vial lids contain a similar tube and filter for proper aeration of the cultures, fresh air is constantly sucked in to the culture head-space as the waste pump pulls air out. In longer experiments, a one-way valve (McMaster-Carr) was installed on the media inlet tube, as any backflow would allow aerosolised bacteria to enter the tube. The valves were sterilised daily with ethanol.

### Experimental Setup

All bioreactor experiments are started by plating a glycerol stock onto LB-agar and growing overnight at 30/37 *^◦^*C for *P. putida*/*E. coli* respectively. A single colony is used to inoculate a 5 mL pre-culture with the same media as the experiment (to reduce effects of metabolism adapting to the media) and incubated shaking at 30/37 *^◦^*C to mid-exponential phase for 4-8 hours (depending on size and age of the colony used). For inoculation, the pre-culture OD is measured, 1 mL of each is centrifuged with an relative centrifugal force (RCF) of 4000 for 4 minutes, and they are resuspended in PBS to reach an OD of 2 to remove pre-culture media and standardise the inoculation concentration. Each bioreactor, which has a culture volume of 20 mL, is inoculated with 100 µL of this PBS suspension for a final OD of 0.01.

All co-culture experiments were conducted by mixing a pair of *P. putida* and *E. coli* monocultures at steady state, in a 50:50 ratio unless stated otherwise. The OD of each culture was first measured with an external spectrometer to ensure they were similar, and they were diluted with M9 otherwise. For increased standardisation, a pair of monocultures was typically mixed into 4 bioreactors: from a 20 mL culture, 16 mL was split into 4 freshly autoclaved/wiped vials, and 1 mL used for measuring OD. Each mixture, which contains a total of 8 mL, is then filled to 20 mL with media. The monocultures were typically grown at an OD of 0.75 (such that the OD after splitting into 4 reactors was around 0.375) without dithering, as they would have different ODs depending on how recently they had been diluted. Before implementing control, the culture is allowed to grow for 2 hours at 33.2 *^◦^*C (the critical temperature) to allow it to reach the OD setpoint and stabilise. While arbitrary references are possible, for long-term experiments we selected a 30:70 *P. putida*:*E. coli* ratio in an attempt to reduce the impact of biofilm formation.

### Offline Validation

Sampling for offline validation was done by pausing the bioreactors and removing 1 mL of culture in a sterile hood. This was centrifuged with an RCF of 14000 for 2 minutes, resuspended in 0.5 mL of PBS, then mixed with 0.5 mL of 50 % glycerol and stored at *−*80 *^◦^*C. For validation, samples were thawed on 4 *^◦^*C metal beads, then washed once and resuspended in 1 mL of 4 *^◦^*C PBS. For flow cytometry, samples were analysed in a FACSAriaIII (BD Biosciences) with a 405 nm laser and 450/40 nm filter for pyoverdine and a 561 nm laser and 610/20 nm filter for RFP. Voltages were as follows: 400 (FSC), 350 (SSC), 650 (561-610/20), and 400 (405-405/40). FSC-A and SSC-A channels were used to gate living cells, and for validating control experiments *P. putida* or *E. coli* monocultures were used as negative/positive controls for fluorescence with the 561 nm laser. Flow cytometry noise was calculated from measurements of 12 replicates of samples taken immediately after mixing (S6 Figure). For plating, samples were diluted serially 8 times in a 96 well plate with PBS. 5 µL of each dilution was then plated onto two different LB-agar plates, one containing 15 µg mL*^−^*^1^ chloramphenicol. Colonies from the least diluted level were counted, where *E. coli* = colonies without Cm - colonies with Cm.

### Biofilm Assay

Biofilms were stained with 0.1 % w/v crystal violet for visualisation and assays. Vials or 96 well plates containing biofilms are emptied of culture by shaking over a waste tray, washed by gently submerging in tap water, then left to dry in a sterile hood for 15 minutes before staining for 10 minutes (22 mL for vials and 220 µL for plates). The stain was poured into a waste tray, and the biofilms were washed thrice by gently submerging them in tap water before being left to dry and stored. For quantification, 95 % ethanol was used to solubilise the crystal violet from the biofilms for 15 minutes, then 125 µL was transferred to a 96 well plate and OD_595_ measured in a platereader. To evaluate the effect of Tween (polysorbate) 20 (S9 Figure) or cellulase (S10 Figure) on biofilm formation, they were added in a range of concentrations to 20 mL of culture in bioreactors or to 200 µL of culture in 96 well plates and grown overnight (bioreactor) or for 24 hours (plates).

## Supporting information

**S1 File. Models utilised in the state estimation.** Mathematical derivation and description of the utilised models and how they were used by the estimator.

**S1 Figure. Calibrating fluorescence readings in different bioreactors.** Each reactor was given a unique identifier and then calibrated (top) by measuring fluorescence (a.u.) of a serial dilution of a filter-sterilised *P. putida* monoculture (removing cells but maintaining fluorescent extracellular pyoverdine). Used excitation/excitation wavelengths of 395/440 nm and diluted with M9 media. These measurements were used to calculate a factor *fl_fac_*(left) and offset *fl_ofs_*(right) for each reactor, where the factor is relative to reactor h.

**S2 Figure. Effect of temperature on media and pyoverdine.** Reactor runs of just fresh media (M9 + CAA, left) and a filtered *P. putida* monoculture (i.e. M9 + CAA + pyoverdine, right). The negligible (*<*30 a.u.) change in fluorescence across a temperature range larger than conditions used in experiments for both plain media and media with pyoverdine indicates that large changes in fluorescence observed in experiments with cells are due to changes in pyoverdine production.

**S3 Figure. Dilution series of *P. putida* from a bioreactor monoculture spotted onto LB-agar plates with chloramphenicol.** Concentration range of 0, 5, 7.5, 10, 12.5, 15, 20 µg mL*^−^*^1^. No significant difference in the growth of *P. putida* observed at any concentration, 15 µg mL*^−^*^1^ used (sufficient to prevent *E. coli* growth-not shown).

**S4 Figure. Number of Colony Foming Units (CFUs) from a dilution series of *P. putida* and *E. coli* from bioreactor monoculture**. Diluted with 20 % intervals using PBS. Technical replicates at each dilution and biological replicates in different reactors vary, possibly due to noise in OD measurement. Flow cytometry, which measures ratios, is unaffected by this type of noise.

**S5 Figure. Freeze-thawing stored samples causes pyoverdine to leak out of cells.** Flow cytometry gating of fresh (left)/freeze-thawed (right) samples from a *P. putida* monoculture, and a freeze-thawed sample from a co-culture (bottom). Cells were first gated by forward and side scatter (not shown), and then by fluorescence intensity for pyoverdine (405 - 450/40-A) or RFP (561 - 610/20-A) wavelengths. Fresh *P. putida* has significant amounts of pyoverdine fluorescence, but almost is lost after freeze-thawing. In the flow cytometer, species are instead distinguished using the RFP’s fluorescence, which persists after a freeze-thaw.

**S6 Figure. Flow cytometer noise.** Histogram of twelve flow cytometer measurements of the same composition. This resulted in an estimated flow cytometry measurement uncertainty with a standard deviation of 5.3 %.

**S7 Figure. OD dithering around setpoint**. The total OD (measured at 600*_nm_*) of the culture dithers around a setpoint of 0.5, growing from 0 until it reaches 0.545, at which point fresh media is pumped in until the OD is diluted down to 0.455. This creates phases of dilution and growth from which we are able to calculate the growth rate of the mono or co-culture). When the temperature changes, the steepness of the curve during the growth period visibly changes as well.

**S8 Figure. Change of the growth rates under a temperature change from** 29 *^◦^*C **to** 36 *^◦^*C. Growth rates of *P. putida* and *E. coli* calculated from monocultures OD data, where dark red background = 36 *^◦^*C, light red background = 29 *^◦^*C. *P. putida* has a laggy response to temperature change in both directions.

**S9 Figure. Polysorbate (Tween) 20 to prevent biofilms. (A)** Range of concentrations of Tween 20 vs OD_595_ of ethanol-solubilised crystal violet used to stain the wells of 96 well plate growing *P. putida*. A higher OD = more biofilm. Compared to the no-Tween control, concentrations *>*0.001 % have significantly less biofilm formation in plates. **(B)** Monoculture of *P. putida* in a reactor vial after 7 hours of growth with 0.01 % Tween 20. A thin biofilm has still formed after 7 hours at the air-water interface, as highlighted in the red box.

**S10 Figure. Cellulase to prevent biofilms. (A)** Range of concentrations of cellulase vs OD_595_ of ethanol-solubilised crystal violet used to stain the wells of 96 well plates growing *E. coli* (top), *P. putida* (middle), or both (bottom). A higher OD = more biofilm. Concentrations above 0.5 mg mL*^−^*^1^ had less *P. putida* biofilm in plates, while cellulase did not have a clear effect on *E. coli* . Co-cultures had more biofilm when cellulase was added, but not in a concentration-dependent way. **(B)** Monocultures grown in a reactor vial overnight with/without 0.5 mg mL*^−^*^1^ cellulase, where depth of purple indicates amount of biofilm. Cellulase caused *E. coli* to form an extremely thick biofilm, while it appeared to slightly inhibit *P. putida* biofilm formation.

**S11 Figure. Effect of temperature on oscillatory behaviour of pyoverdine production.** Fluorescence is affected by the dithering behaviour of the culture (spiky increases/drops as the culture grows/is diluted)-this is not seen in the Fig 4 D/E, which were grown without dithering (i.e. very small dilutions whenever OD exceeded the setpoint, leading to an almost constant OD). Aside from the spikes, oscillations are also visible at 26 *^◦^*C. The amplitude of the oscillations appears to be related to temperature, decreasing at warmer temperatures.

**S12 Figure. Effect of arginine with complex amino acid mixtures on oscillatory behaviour of pyoverdine production.** *P. putida* monocultures grown with 0.2 % CAA and 1 g L*^−^*^1^ arginine (top) still has dampened oscillatory behaviour at 29 *^◦^*C. When grown with 0.2 % CAA but no arginine, and then spiked with 1 g L*^−^*^1^ arginine at around time = 13 hours (bottom), pyoverdine production drops briefly before rising slowly.

**S13 Figure. Effect of arginine with simple amino acid mixtures on oscillatory behaviour of pyoverdine production.** *P. putida* monocultures grown with 3 g L*^−^*^1^ serine and 0.5 g L*^−^*^1^ glutamine, using values suggested from Maser et al. (2020) [38] along with a range of arginine concentrations. Arginine has a clear effect, with lower concentrations appearing to dampen oscillations but cause a peak in pyoverdine before settling. Excess arginine creates oscillations that did not dampen after 15 hours.

**S14 Figure. Effect of arginine and different synthetic amino acid mixtures on oscillatory behaviour of pyoverdine production.** *P. putida* monocultures grown with double dropout Complete Synthetic Mixtures (- arginine and -tryptophan, both thought to influence pyoverdine production [28]) from Fromedium (SKU: DCS0441). CSM1 (A, B) is supplemented with 100 mg L*^−^*^1^ serine, while CSM2 (C, D) is supplemented with 100 mg L*^−^*^1^ serine and glutamine. Runs B and D were spiked with 100 mg L*^−^*^1^ arginine at around time = 22 hours, which caused production to briefly drop before rising.

**S15 Figure. Effect of CAA concentration on growth rate.** Growth rate of monocultures cultured at a range of CAA concentrations. Because *E. coli* is much more affected by decreasing [CAA] than *P. putida*, the critical temperature is increased and there is a smaller temperature window to select for *E. coli* . The smaller difference in relative growth rate means that the system would respond slower to control inputs.

**S16 Figure. Vials from the long-term co-culture control experiments, t=5 days.** Vials are stained with crystal violet, depth of purple represents amount of biofilm. Open-loop technical replicates had no controller, but the media temperature was set to the critical temperature. Closed-loop technical replicates uses the controller. Closed-loop vials have significantly less biofilm, presumably because the *P. putida* population was maintained at around 30 %.

**S17 Figure. Additional replicates of long-term control.** Similar trends are observed to replicates from Fig 7, where open-loop control maintains the composition around the setpoint for a day, but then experiences large fluctuations over the rest of the week. The closed-loop control is still around the setpoint by the end of the experiment, though the *P. putida* abundance rose to around 50 % on the second last day.

## Acknowledgements

The authors thank Robert Hedley and Vasiliki Tsioligka for providing technical assistance with flow cytometry at The Don Mason Facility of Flow Cytometry, Sir William Dunn School of Pathology, University of Oxford. Professor Tim Tolker Nielsen and Dr Morten Rybtke at Costerton Biofilm Centre at the University of Copenhagen are greatly thanked for their kind gift of *P. putida* OUS82 and derivative strains.

## References

1. Shin J, Liao S, Kuanyshev N, Xin Y, Kim C, Lu T, et al. Compositional and temporal division of labor modulates mixed sugar fermentation by an engineered yeast consortium. Nature Communications. 2024;15(1):781. doi:10.1038/s41467-024-45011-w.

2. Tsoi R, Wu F, Zhang C, Bewick S, Karig D, You L. Metabolic division of labor in microbial systems. Proceedings of the National Academy of Sciences. 2018;115(10):2526–2531. doi:10.1073/pnas.1716888115.

3. Gasparek M, Steel H, Papachristodoulou A. Deciphering mechanisms of production of natural compounds using inducer-producer microbial consortia. Biotechnology Advances. 2023;64:108117. doi:10.1016/j.biotechadv.2023.108117.

4. Stenuit B, Agathos SN. Deciphering microbial community robustness through synthetic ecology and molecular systems synecology. Current Opinion in Biotechnology. 2015;33:305–317. doi:10.1016/j.copbio.2015.03.012.

5. Xia T, Eiteman MA, Altman E. Simultaneous utilization of glucose, xylose and arabinose in the presence of acetate by a consortium of Escherichia coli strains. Microbial Cell Factories. 2012;11(1):77. doi:10.1186/1475-2859-11-77.

6. Hardin G. The Competitive Exclusion Principle. Science. 1960;131(3409):1292–1297. doi:10.1126/science.131.3409.1292.

7. Dinh CV, Chen X, Prather KLJ. Development of a Quorum-Sensing Based Circuit for Control of Coculture Population Composition in a Naringenin Production System. ACS Synthetic Biology. 2020;9(3):590–597. doi:10.1021/acssynbio.9b00451.

8. Salma A, Abdallah R, Fourcade F, Amrane A, Djelal H. A New Approach to Produce Succinic Acid Through a Co-Culture System. Applied Biochemistry and Biotechnology. 2021;193(9):2872–2892. doi:10.1007/s12010-021-03572-2.

9. Mee MT, Collins JJ, Church GM, Wang HH. Syntrophic exchange in synthetic microbial communities. Proceedings of the National Academy of Sciences. 2014;111(20):E2149–E2156. doi:10.1073/pnas.1405641111.

10. Miano A, Liao MJ, Hasty J. Inducible cell-to-cell signaling for tunable dynamics in microbial communities. Nature Communications. 2020;11(1):1193. doi:10.1038/s41467-020-15056-8.

11. Boo A, Mehta H, Amaro RL, Stan GB. Host-aware RNA-based control of synthetic microbial consortia; 2023. Available from: https://www.biorxiv.org/content/10.1101/2023.05.15.540816v1.

12. McCarty NS, Ledesma-Amaro R. Synthetic Biology Tools to Engineer Microbial Communities for Biotechnology. Trends in Biotechnology. 2019;37(2):181–197. doi:10.1016/J.TIBTECH.2018.11.002.

13. Caringella G, Bandiera L, Menolascina F. Recent advances, opportunities and challenges in cybergenetic identification and control of biomolecular networks. Current Opinion in Biotechnology. 2023;80:102893. doi:10.1016/j.copbio.2023.102893.

14. Bertaux F, Sosa-Carrillo S, Gross V, Fraisse A, Aditya C, Furstenheim M, et al. Enhancing bioreactor arrays for automated measurements and reactive control with ReacSight. Nature Communications. 2022;13(1):3363. doi:10.1038/s41467-022-31033-9.

15. Aditya C, Bertaux F, Batt G, Ruess J. A light tunable differentiation system for the creation and control of consortia in yeast. Nature Communications. 2021;12(1):5829. doi:10.1038/s41467-021-26129-7.

16. Gutíerrez J, Kumar S, Khammash M. Dynamic cybergenetic control of bacterial co-culture composition via optogenetic feedback. Nature Communications. 2022;13(1):4808. doi:10.1038/s41467-022-32392-z.

17. Lee TA, Steel H. Cybergenetic control of microbial community composition. Frontiers in Bioengineering and Biotechnology. 2022;10.

18. Kusuda M, Shimizu H, Toya Y. Reactor control system in bacterial co-culture based on fluorescent proteins using an Arduino-based home-made device. Biotechnology Journal. 2021;16(12):2100169. doi:10.1002/biot.202100169.

19. Khammash M, Di Bernardo M, Di Bernardo D. Cybergenetics: Theory and Methods for Genetic Control System. In: 2019 IEEE 58th Conference on Decision and Control (CDC); 2019. p. 916–926.

20. Carrasco-Ĺopez C, Garćıa-Echauri SA, Kichuk T, Avalos JL. Optogenetics and biosensors set the stage for metabolic cybergenetics. Current Opinion in Biotechnology. 2020;65:296–309. doi:10.1016/j.copbio.2020.07.012.

21. Steel H, Habgood R, Kelly C, Papachristodoulou A. In situ characterisation and manipulation of biological systems with Chi.Bio. PLOS Biology. 2020;18(7):e3000794. doi:10.1371/JOURNAL.PBIO.3000794.

22. Ainelo H, Lahesaare A, Teppo A, Kivisaar M, Teras R. The promoter region of lapA and its transcriptional regulation by Fis in Pseudomonas putida. PLoS ONE. 2017;12(9):e0185482. doi:10.1371/journal.pone.0185482.

23. Gjermansen M, Nilsson M, Yang L, Tolker-Nielsen T. Characterization of starvation-induced dispersion in Pseudomonas putida biofilms: genetic elements and molecular mechanisms. Molecular Microbiology. 2010;75(4):815–826. doi:10.1111/j.1365-2958.2009.06793.x.

24. Hoegy F, Mislin GLA, Schalk IJ. Pyoverdine and Pyochelin Measurements. In: Filloux A, Ramos JL, editors. Pseudomonas Methods and Protocols. New York, NY: Springer; 2014. p. 293–301. Available from: 10.1007/978-1-4939-0473-0_24.

25. Krieger AG, Zhang J, Lin XN. Temperature regulation as a tool to program synthetic microbial community composition. Biotechnology and Bioengineering. 2021;118(3):1381–1392. doi:10.1002/bit.27662.

26. Sloup RE, Cieza RJ, Needle DB, Abramovitch RB, Torres AG, Waters CM. Polysorbates prevent biofilm formation and pathogenesis of Escherichia coli O104:H4. Biofouling. 2016;32(9):1131. doi:10.1080/08927014.2016.1230849.

27. Kamali E, Jamali A, Izanloo A, Ardebili A. In vitro activities of cellulase and ceftazidime, alone and in combination against Pseudomonas aeruginosa biofilms. BMC Microbiology. 2021;21:347. doi:10.1186/s12866-021-02411-y.

28. Barrientos-Moreno L, Molina-Henares MA, Pastor-Garćıa M, Ramos-Gonźalez MI, Espinosa-Urgel M. Arginine Biosynthesis Modulates Pyoverdine Production and Release in Pseudomonas putida as Part of the Mechanism of Adaptation to Oxidative Stress. Journal of Bacteriology. 2019;201(22):e00454–19. doi:10.1128/JB.00454-19.

29. Fleurial M, Sacchelli L, Yabo AG. State estimation in alcoholic fermentation models: a case-study in wine-making conditions; 2023. Available from: https://hal.science/hal-04269299.

30. Hellmann S, Wilms T, Streif S, Weinrich S. Comparison of Unscented Kalman Filter Design for Agricultural Anaerobic Digestion Model; 2024. Available from: http://arxiv.org/abs/2310.15958.

31. Asswad R, Cinquemani E, Gouźe JL. Kalman-based approaches for online estimation of bioreactor dynamics from fluorescent reporter measurements; 2024. Available from: http://arxiv.org/abs/2404.16649.

32. Åström KJ, Murray R. Feedback Systems: An Introduction for Scientists and Engineers, Second Edition. Princeton University Press; 2021.

33. Toll-Riera M, Olombrada M, Castro-Giner F, Wagner A. A limit on the evolutionary rescue of an Antarctic bacterium from rising temperatures. Science Advances. 2022;8(28):eabk3511. doi:10.1126/sciadv.abk3511.

34. Lehmann M, Prohaska C, Zeldes B, Poehlein A, Daniel R, Basen M. Adaptive laboratory evolution of a thermophile toward a reduced growth temperature optimum. Frontiers in Microbiology. 2023;14. doi:10.3389/fmicb.2023.1265216.

35. Weinstock MT, Hesek ED, Wilson CM, Gibson DG. Vibrio natriegens as a fast-growing host for molecular biology. Nature Methods. 2016;13(10):849–851. doi:10.1038/nmeth.3970.

36. Bechhofer DH, Stasinopoulos SJ. tetA(L) Mutants of a Tetracycline-Sensitive Strain of Bacillus subtilis with the Polynucleotide Phosphorylase Gene Deleted. Journal of Bacteriology. 1998;180(13):3470–3473.

37. Miyashiro T, Ruby EG. Shedding light on bioluminescence regulation in Vibrio fischeri. Molecular Microbiology. 2012;84(5):795–806. doi:10.1111/j.1365-2958.2012.08065.x.

38. Maser A, Peebo K, Vilu R, Nahku R. Amino acids are key substrates to *Escherichia coli* BW25113 for achieving high specific growth rate. Research in Microbiology. 2020;171(5):185–193. doi:10.1016/j.resmic.2020.02.001.

