## Supplementary figures and images for "Cybernetic control of a natural microbial co-culture"

### S1 Figure

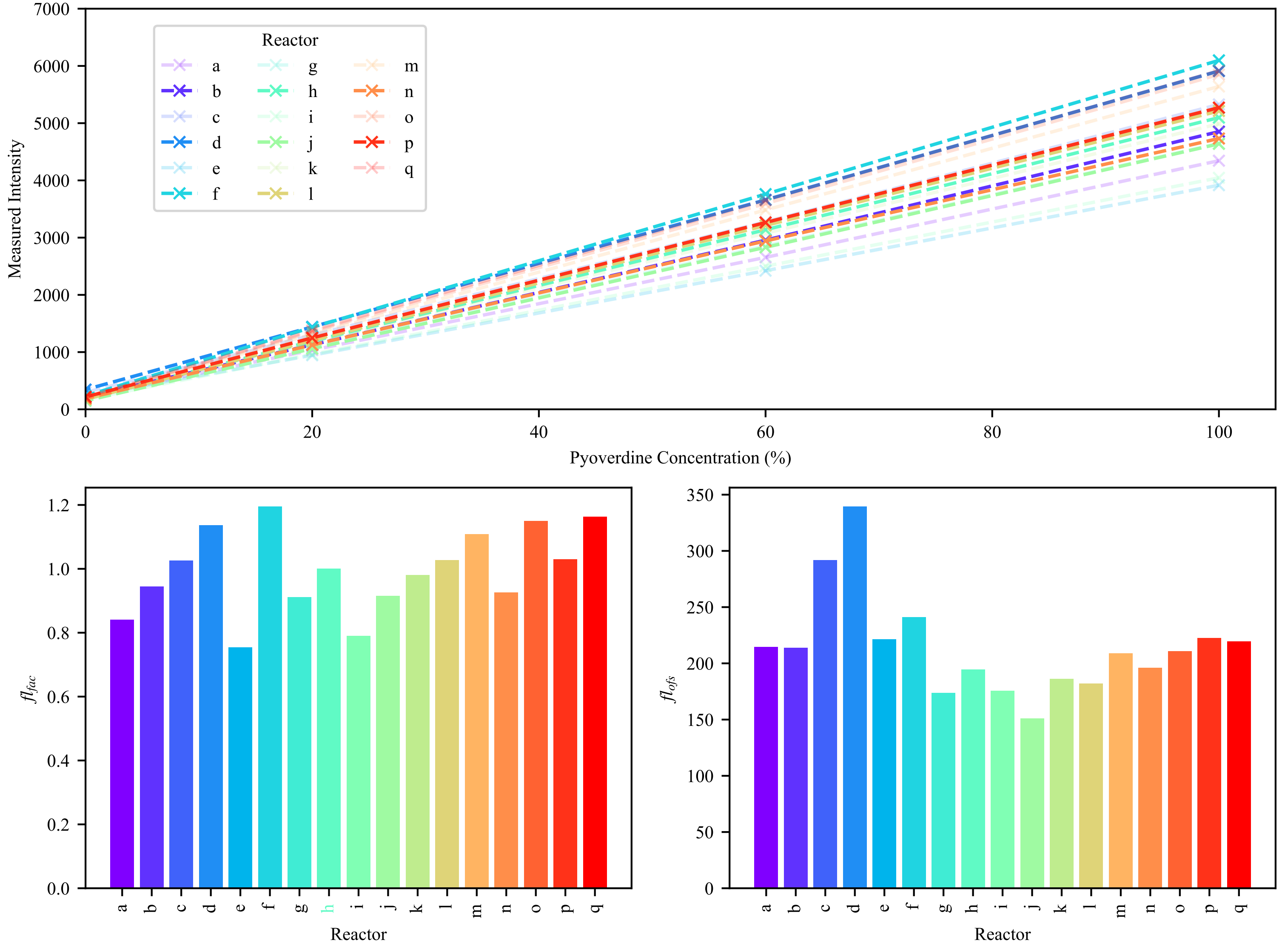

### S2 Figure

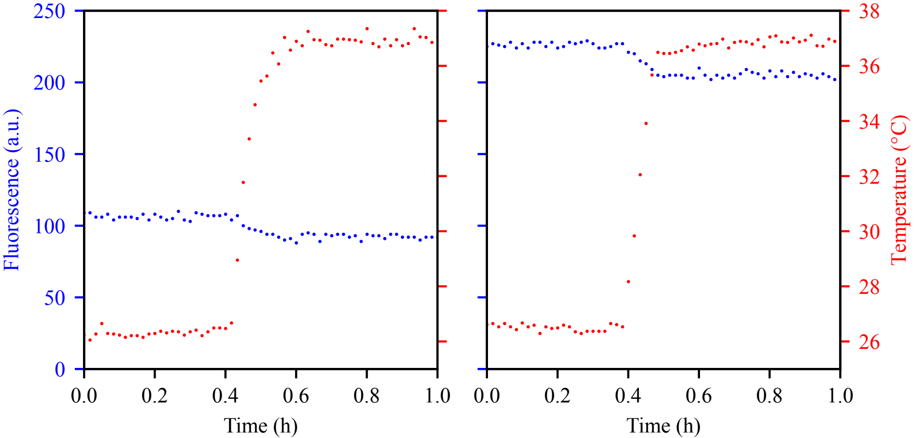

### S3 Figure

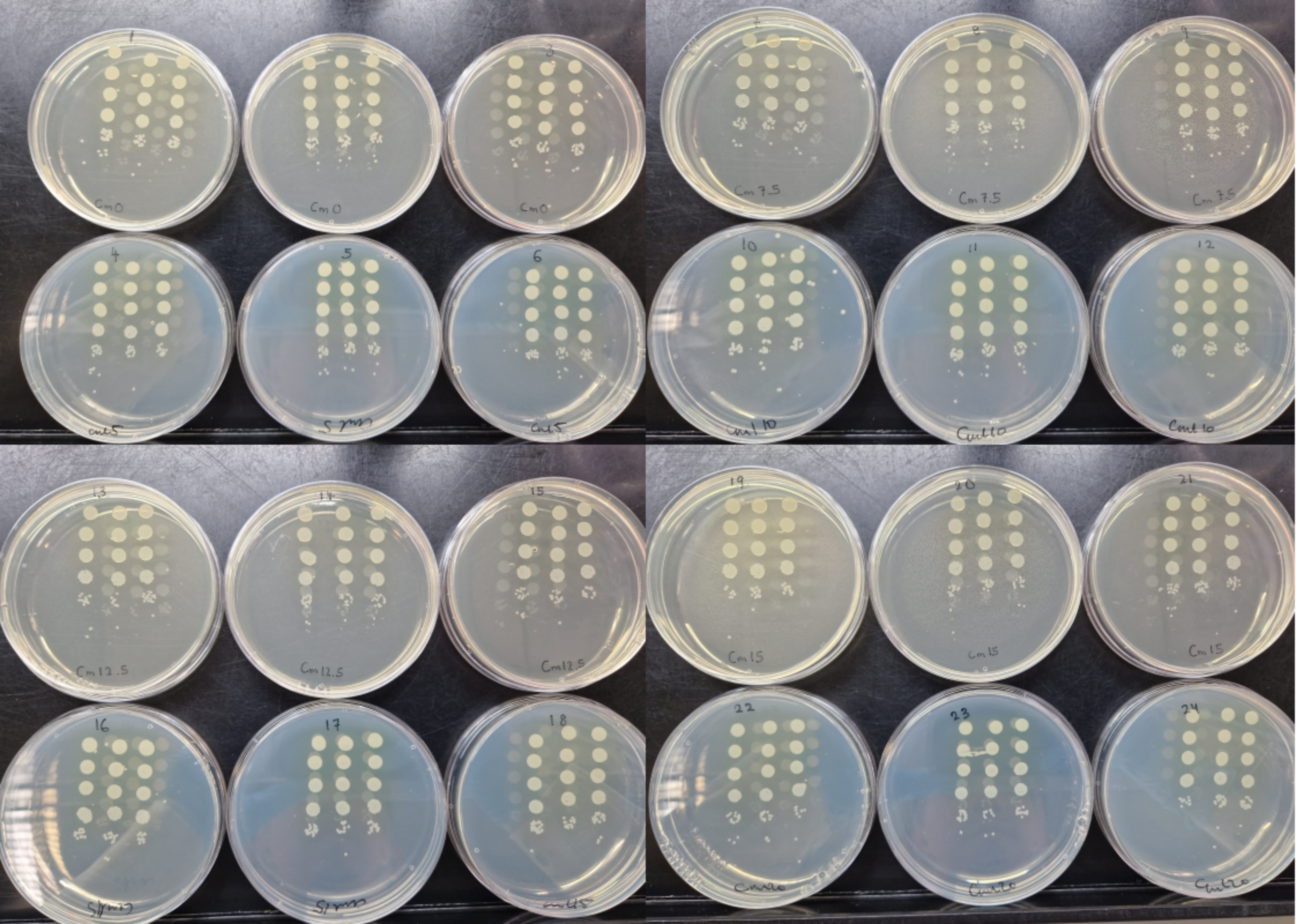

### S4 Figure

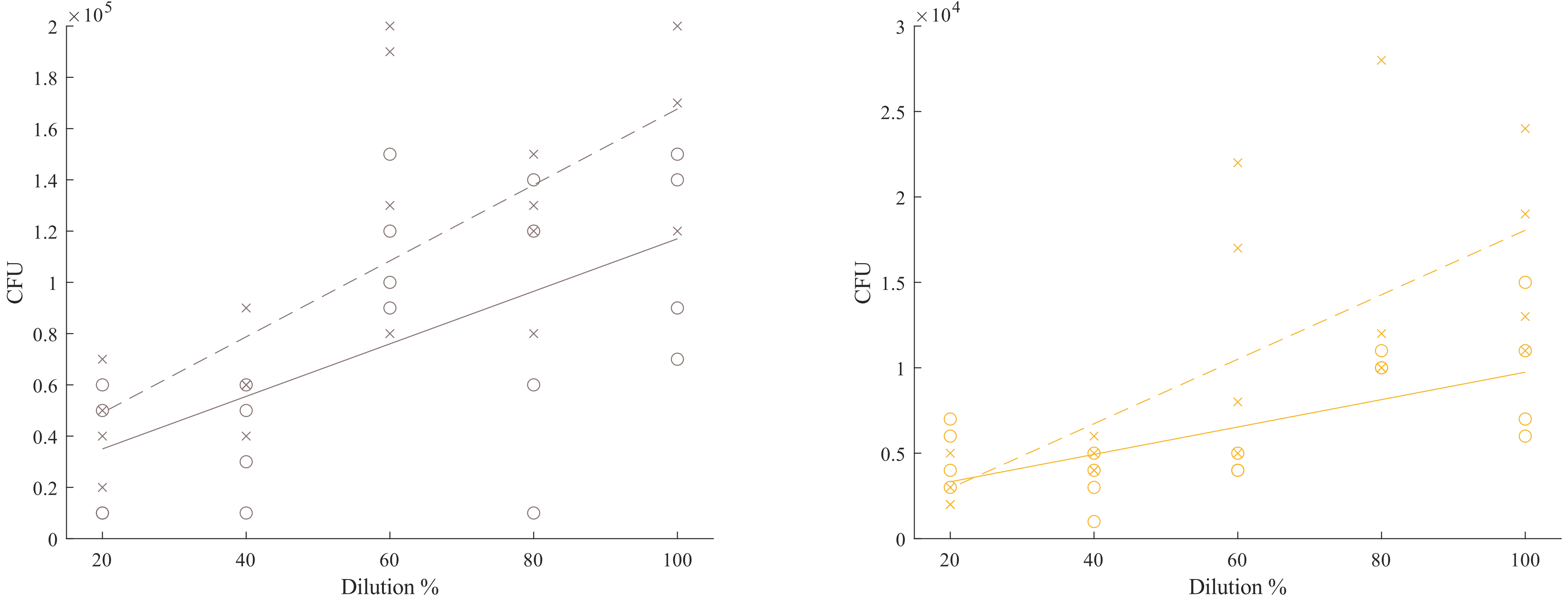

### S5 Figure

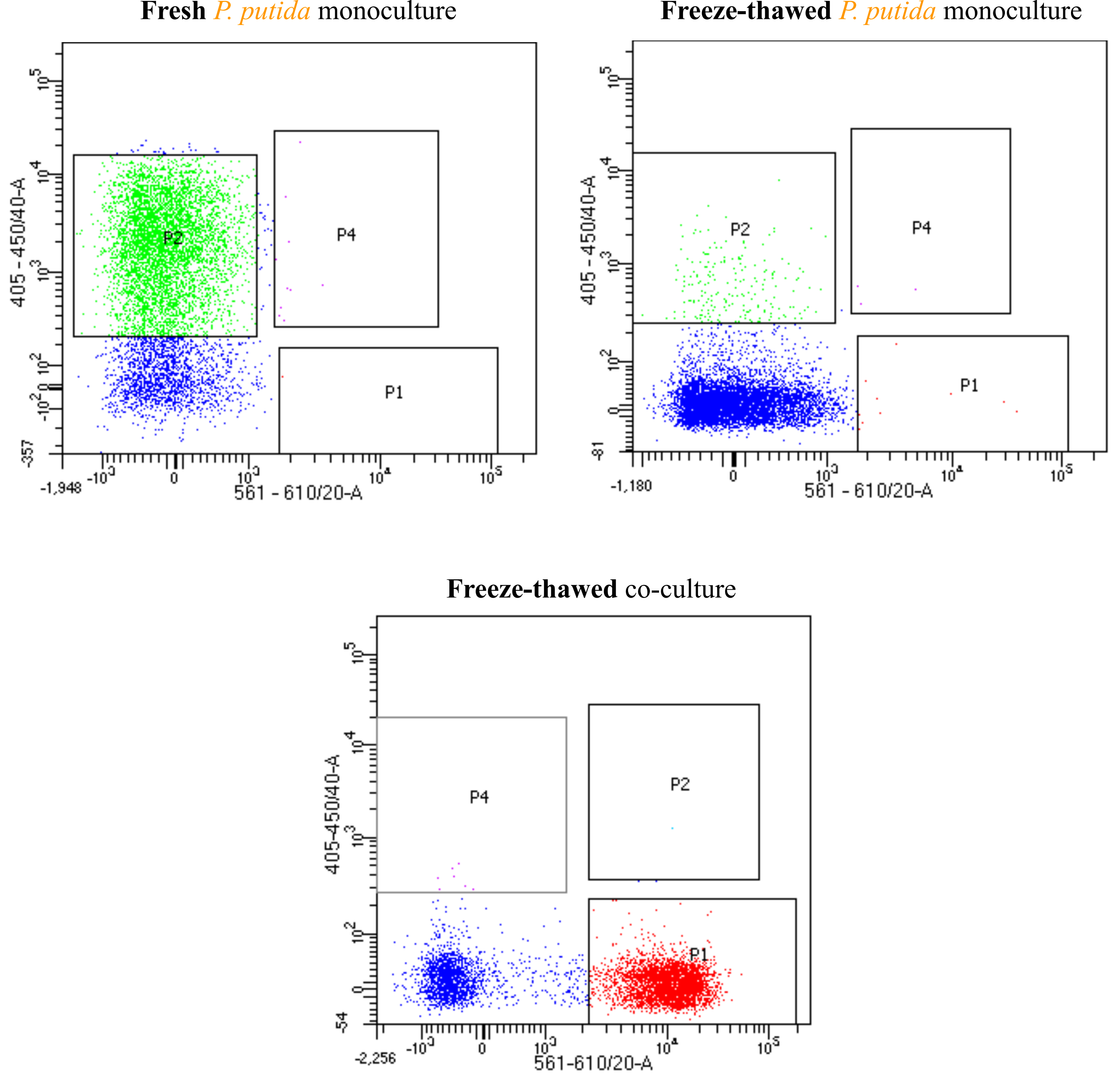

### S6 Figure

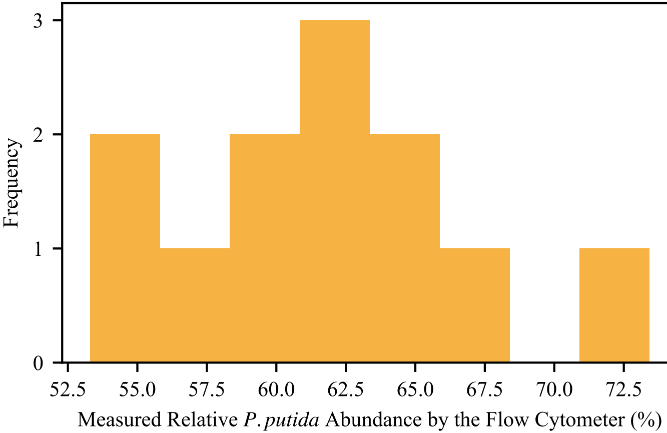

### S7 Figure

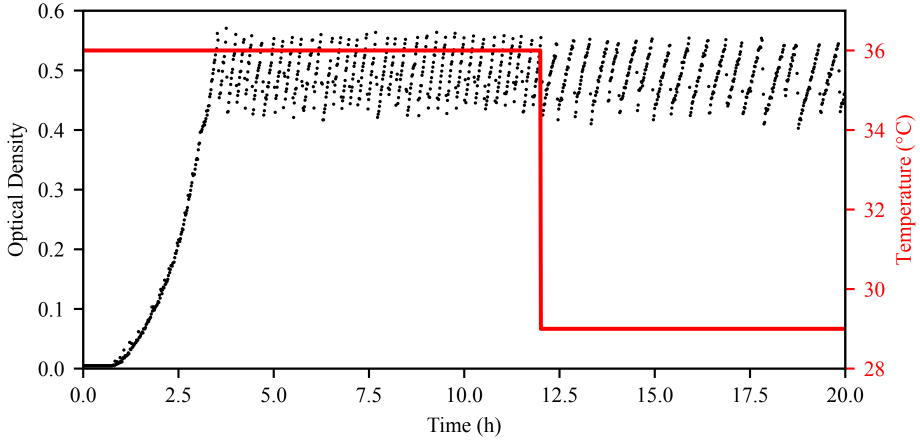

### S8 Figure

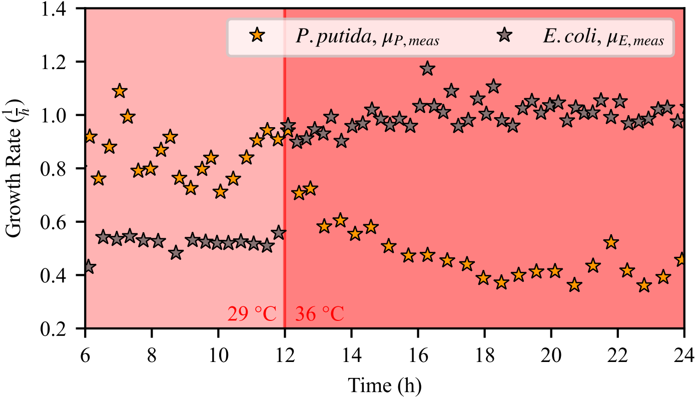

### S9 Figure

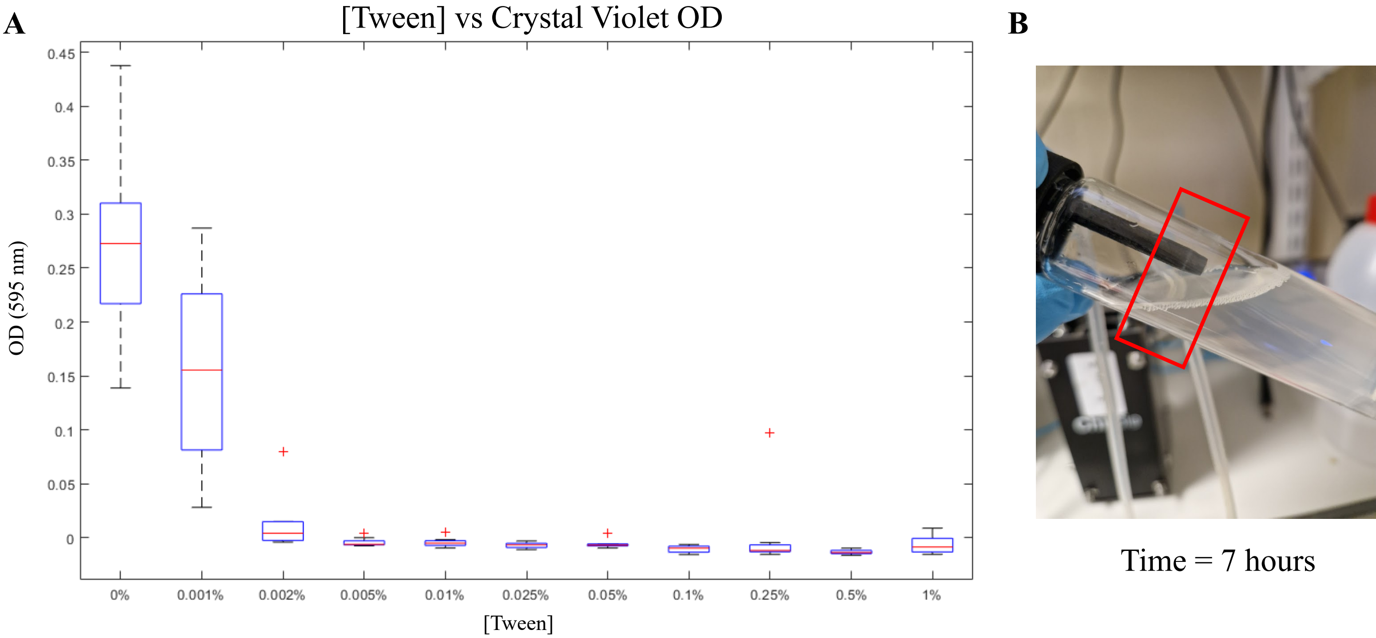

### S10 Figure

**A**

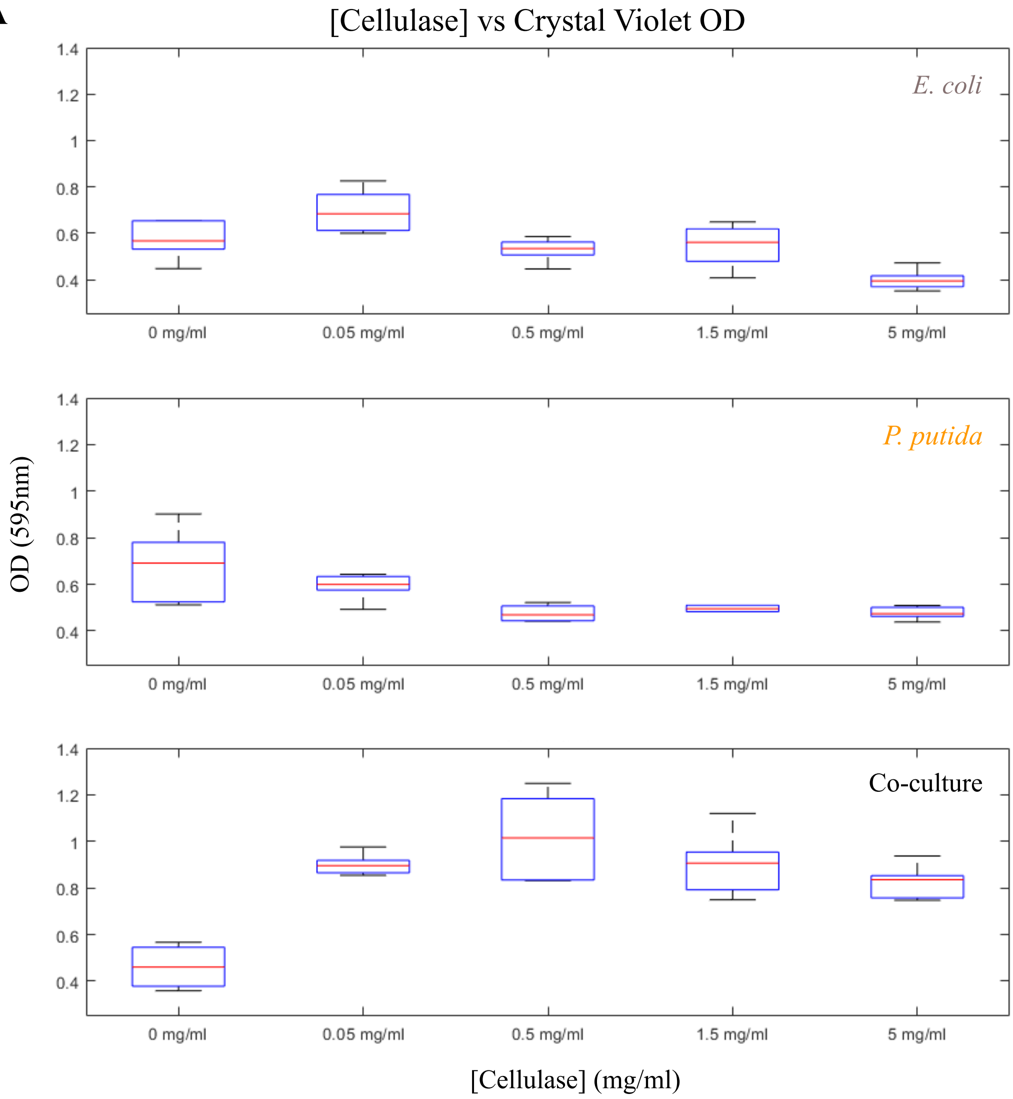


**B**

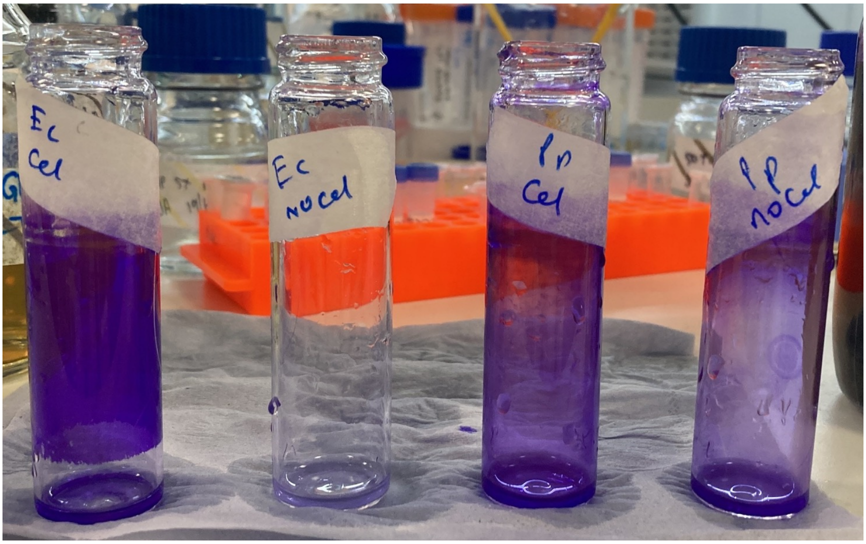


| | | | |
|---|---|---|---|
| *E. coli* | *E. coli* | *P. putida* | *P. putida* |
| + cellulase | - cellulase | + cellulase | - cellulase |

### S11 Figure

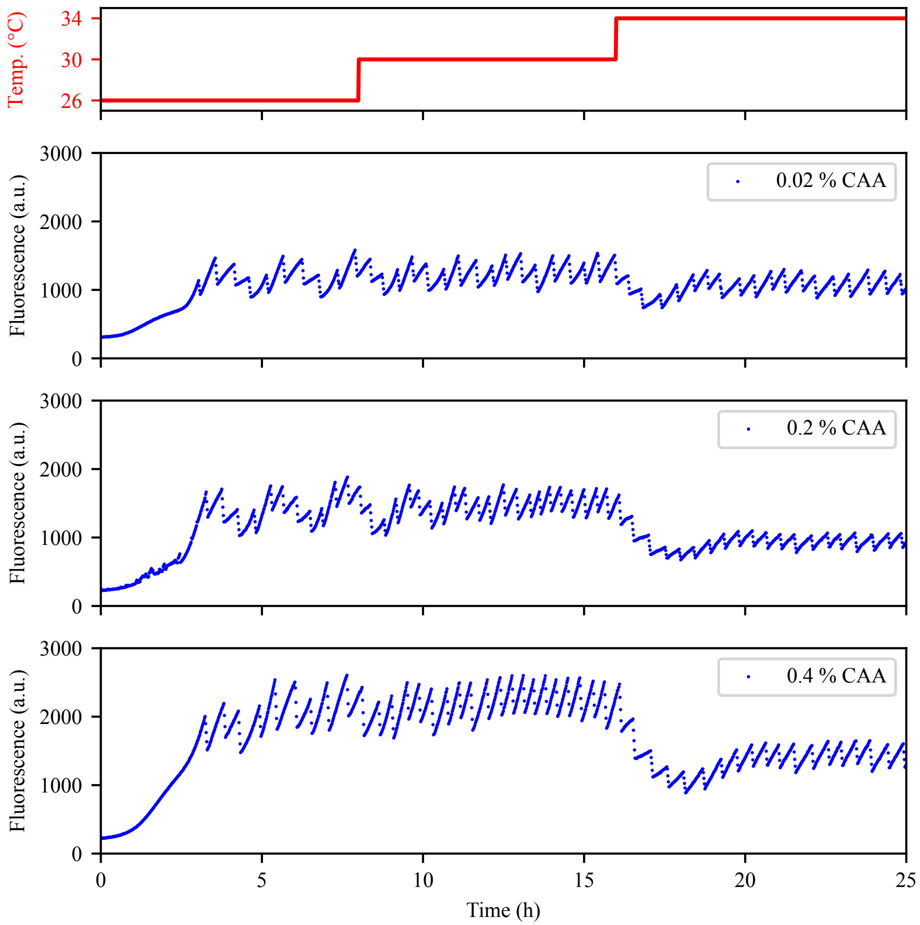

### S12 Figure

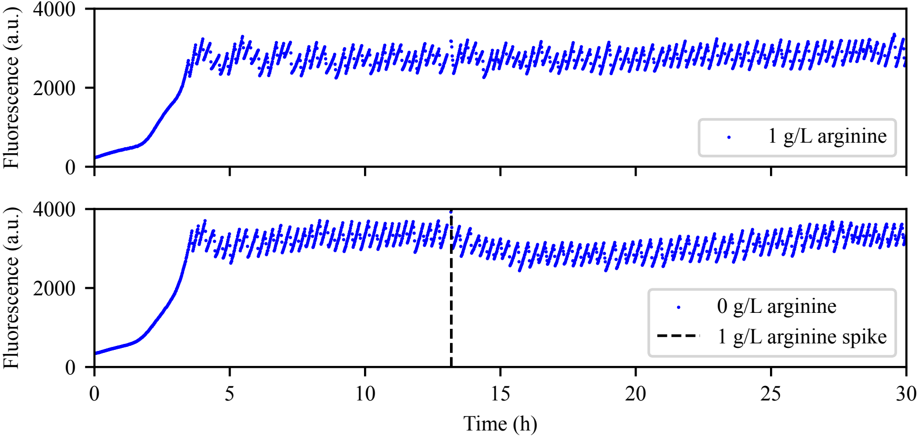

### S13 Figure

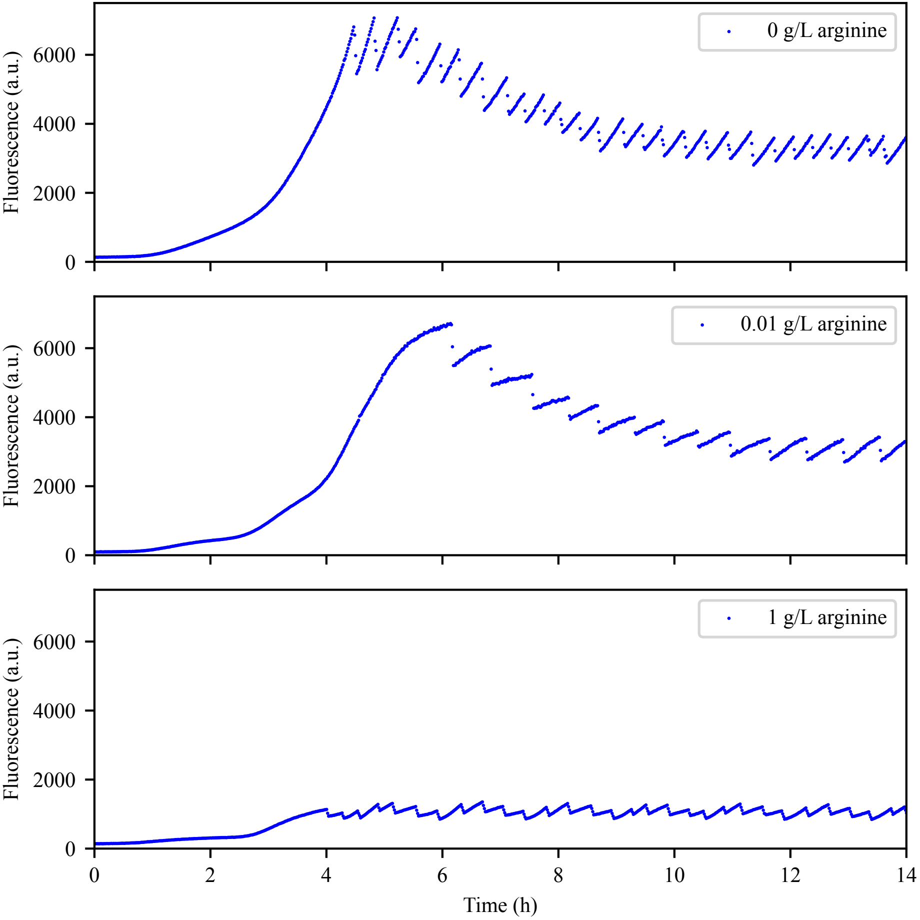

### S14 Figure

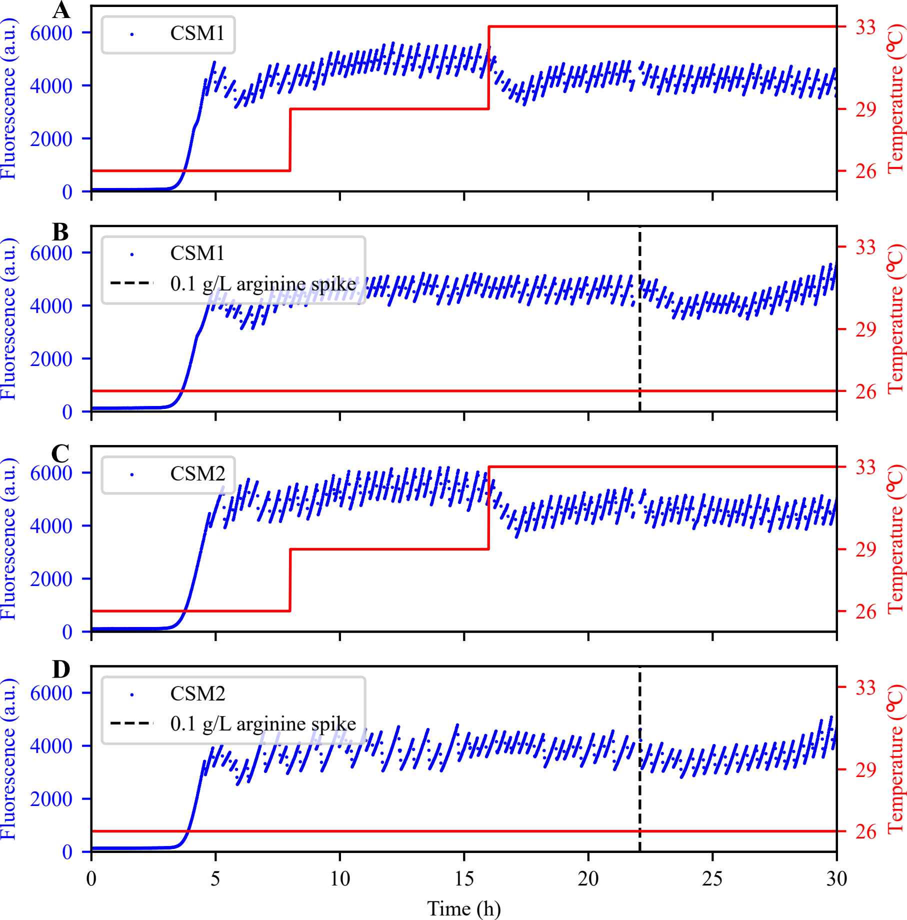

### S15 Figure

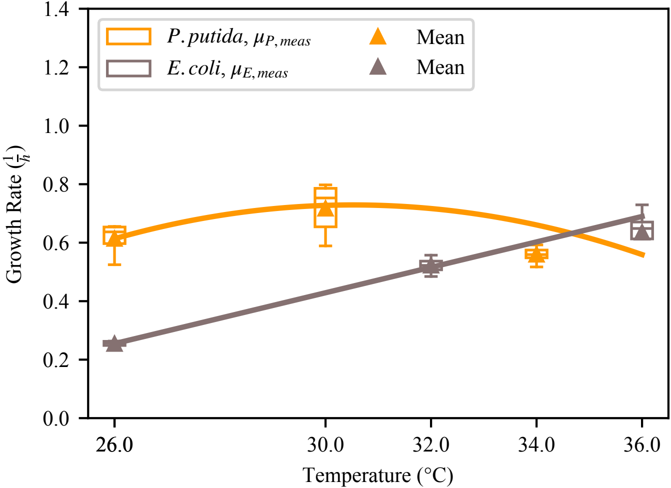

### S17 Figure

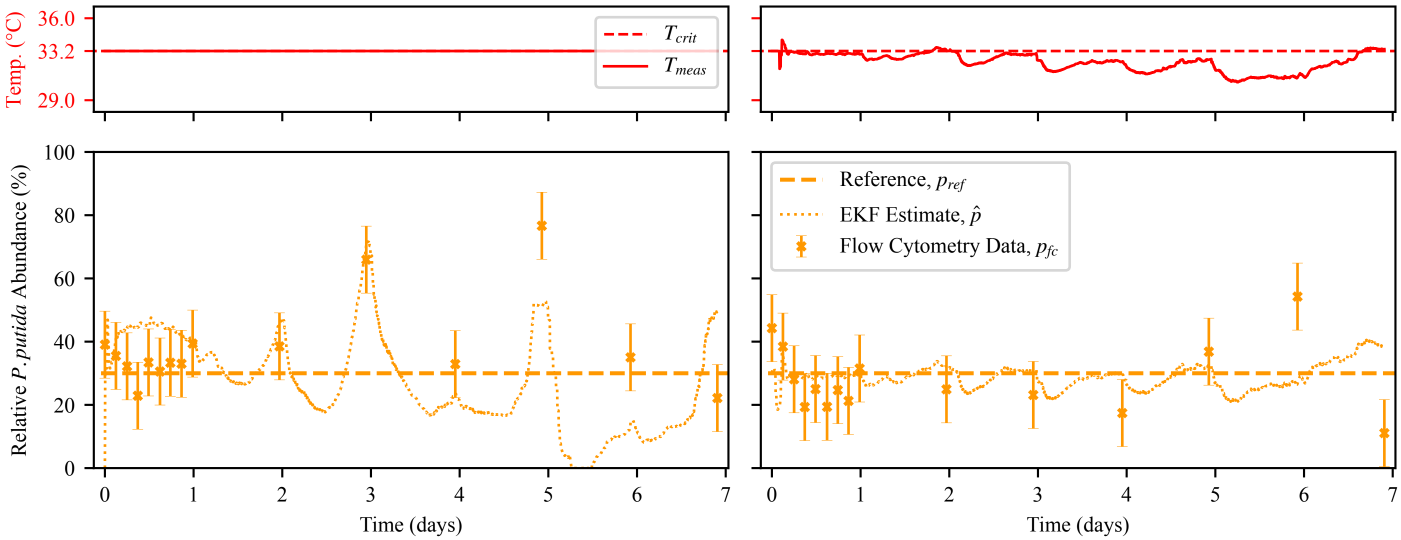
