## Supplementary material for "Cybernetic control of a natural microbial co-culture": S16 Figure

Vials from Long-term Control Experiment, t = 5 days

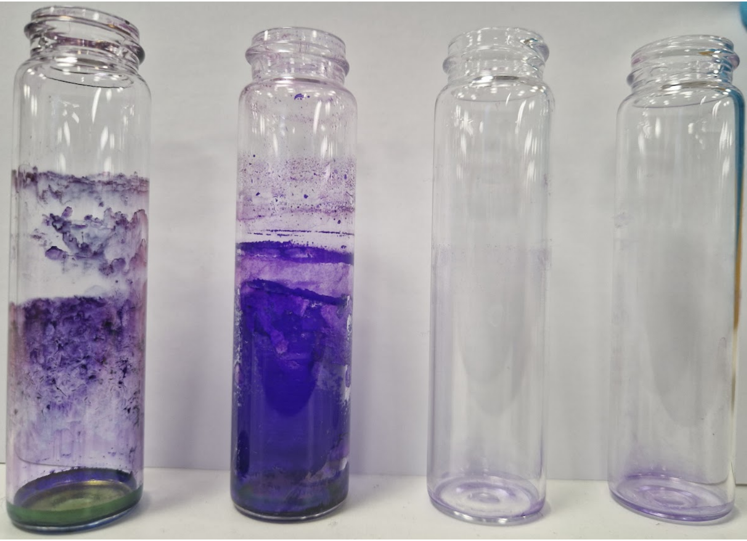

Open Loop tech rep 1 | Open Loop tech rep 2 | Closed Loop tech rep 2 | Closed Loop tech rep 2
