## Supplementary material for "Cybernetic control of a natural microbial co-culture": S1 File

#### S1 Models utilised in the state estimation

##### S1.1 System Model

The system model governs the dynamics of three key components, i.e. the $E(t)$ represents the optical density (OD) of *Echerichia coli* (*E. coli*), $P(t)$ the OD of *Pseudomonas putida* (*P. putida*), and $F[k]$ the fluorescence level of pyoverdine. To better observe bacteria growth and pyoverdine production, and hence derive a more accurate composition estimate, a zigzag pattern in OD was created (already described in [1]). This resulted in a growth period, without any dilution and rising OD, and a dilution period, where the OD got reduced until a lower threshold was reached. As such, the system model was split into growth and dilution models to account for the different phases of the reactor and thus co-culture dynamics.

###### S1.1.1 Growth Period

It was assumed that substrates in the media are at high abundance and their change of concentration is negligible. At low OD, bacterial growth can be modeled by:

$$\begin{aligned} \dot{P}(t) &= \mu_P(t)P(t) \\ \dot{E}(t) &= \mu_E(t)E(t) \end{aligned} \tag{M.1}$$

Where the temperature-dependent and hence time-dependent growth rate $\mu(t)$ describes how fast the OD of the individual culture increases at time $t$. Similarly to the state variables $E(t)$ and $P(t)$, the growth rates relate to the change of bacteria OD over time and do not describe the cell's individual growth. Euler Forward discretization of the ordinary differential equations (ODEs) in Equation (M.1) with sampling time $t_s$ result in:

$$\begin{aligned} P[k+1] &= (1 + t_s(\mu_P[k] + \nu_P[k]))P[k] \\ E[k+1] &= (1 + t_s(\mu_E[k] + \nu_E[k]))E[k] \end{aligned} \tag{M.2}$$

with process noise $\nu_P[k]$ and $\nu_E[k]$ at cycle $k$. Assuming that the production of pyoverdine only linearly depends on the present amount of *P. putida* and production rate $\mu_F[k]$, pyoverdine fluorescence $F[k]$ with process noise $\nu_F[k]$ can be modelled as follows:

$$F[k+1] = F[k] + (t_s(\mu_F[k] + \nu_F[k]))P[k] \tag{M.3}$$

The process noise was multiplicative to model the increased absolute uncertainty at higher bacteria abundance. Intuitively, it described the uncertainty of the defined growth and production rates.

Temperature influence on growth and production rates were determined in separate monocultures through OD and fluorescence measurements between dilutions. The detailed computation of the rates from the measurements is given in subsection S1.4 and subsection S1.5. The measured rates converged to a steady state for each temperature after passing a transient phase (Fig M1.1). The respective rates at steady state for the relevant temperatures are displayed in detail (Fig 2B and Fig 3A). By fitting polynomial models, we obtain the steady-state bacterial growth rates $\mu_{P,ss}[T]$, $\mu_{E,ss}[T]$, and $\mu_{F,ss}[T]$ as a function of the temperature $T$. To accurately describe the transient phase of the *P. putida* and

*E. coli* growth rates we add dynamics to the growth and production rates with a first-order infinite impulse response (IIR) filter:

$$\begin{aligned} \mu_P[k+1] &= \mu_P[k] + \alpha_P(\mu_{P,ss}[T] - \mu_P[k]) \\ \mu_E[k+1] &= \mu_E[k] + \alpha_E(\mu_{E,ss}[T] - \mu_E[k]) \end{aligned} \tag{M.4}$$

The pyoverdine production rate displayed damped oscillations after a temperature change. These were modelled by extending the IIR filter by a lag of $k_d$ cycles:

$$\mu_F[k+1] = \mu_F[k] + \alpha_F(\mu_{F,ss}[T] - \mu_F[k-k_d]) \tag{M.5}$$

Due to the observed variation of these oscillations in magnitude and period, a conservative model, that did not match the observed data perfectly, was chosen to limit the risk of overfitting (Fig M1.1B).

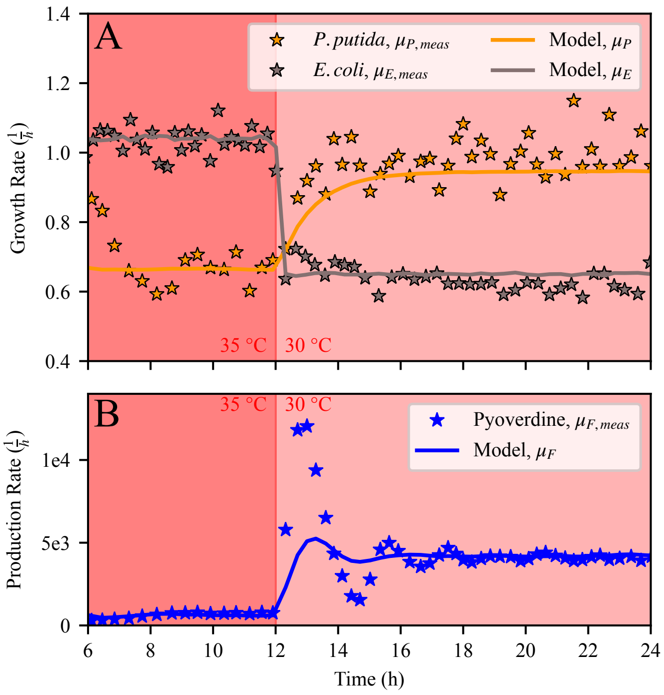


Figure M1.1: Measured rates of bacterial growth ($\mu_{P,meas}$ and $\mu_{E,meas}$) (A) and Pyoverdine production ($\mu_{F,meas}$) (B) for the temperature change from 35 °C to 30 °C. The rates were obtained from the curvature of OD and fluorescent measurements between dilutions. Modelling the rate dynamics over the different temperatures resulted in $\mu_P$, $\mu_E$, and $\mu_F$ (solid lines).

##### S1.1.2 Dilution Period

In the chosen reactor configuration, OD dropped on average by $r_{dil}$ per cycle during dilution. Hence, each quantity decreased as described in the dilution model with multiplicative process noise $\nu_{P,dil}$, $\nu_{E,dil}$, and $\nu_{F,dil}$:

$$\begin{aligned} P[k+1] &= P[k](1 - \frac{r_{dil}}{E[k]+P[k]} + \nu_{P,dil}) \\ E[k+1] &= E[k](1 - \frac{r_{dil}}{E[k]+P[k]} + \nu_{E,dil}) \\ F[k+1] &= F[k](1 - \frac{r_{dil}}{E[k]+P[k]} + \nu_{F,dil}) \end{aligned} \tag{M.6}$$

In the dilution period, these equations were added to the growth period model. All introduced process noises

$$\boldsymbol{\nu} = \begin{pmatrix} \nu_P \\ \nu_E \\ \nu_F \end{pmatrix} \quad \boldsymbol{\nu}_{dil} = \begin{pmatrix} \nu_{P,dil} \\ \nu_{E,dil} \\ \nu_{F,dil} \end{pmatrix}$$

were modelled to be zero-mean and mutually independent, i.e.:

$$\mathbb{E}(\boldsymbol{\nu}) = 0 \qquad \mathbb{E}(\boldsymbol{\nu}_{dil}) = 0$$

$$\mathrm{Cov}(\boldsymbol{\nu}) = \mathbf{Q} = \begin{pmatrix} \sigma_P^2 & 0 & 0 \\ 0 & \sigma_E^2 & 0 \\ 0 & 0 & \sigma_F^2 \end{pmatrix} \qquad \mathrm{Cov}(\boldsymbol{\nu}_{dil}) = \mathbf{Q}_{dil} = \begin{pmatrix} \sigma_{P,dil}^2 & 0 & 0 \\ 0 & \sigma_{E,dil}^2 & 0 \\ 0 & 0 & \sigma_{F,dil}^2 \end{pmatrix}$$

#### S1.2 Measurement Model

The measurement models are important for the composition estimation and the simulation. The Chi.Bio measured the culture's temperature, OD, and fluorescence once per cycle. While the extended Kalman filter (EKF) included the measured temperature directly to derive growth and production rates, it required a measurement model for the latter two rates. OD was treated as a straightforward measurement assumed to be directly proportional to the combined bacteria OD. After sensor calibration, it was modelled with measurement noise $\omega_{od}$, as follows:

$$od[k] = P[k] + E[k] + \omega_{od} \tag{M.7}$$

Bulk fluorescence measurements, $fl[k]$, were acquired by exciting the consortia at 395 nm with light intensity $I_{ex}$ and measuring fluorescence at an emission band around 440 nm. Broad LED emission spectra, imperfect light filters, and light scattering caused the measurement to be affected by bacteria abundance, i.e. OD. This relationship was assumed to be linear, with proportionality constant $c_{od}$. This resulted together with measurement noise $\omega_{fl}$ in:

$$fl[k] = F[k] + c_{od} od[k] + \omega_{fl} \tag{M.8}$$

The parameter $c_{od}$ was obtained from fluorescence measurements under changing OD and in the absence of pyoverdine ($F[k] = 0$). The introduced measurement noise

$$\boldsymbol{\omega} = \begin{pmatrix} \omega_{od} \\ \omega_{fl} \end{pmatrix}$$

was modelled to be zero-mean and mutually independent, i.e.:

$$\mathbb{E}(\boldsymbol{\omega}) = 0 \quad \mathrm{Cov}(\boldsymbol{\omega}) = \begin{pmatrix} \sigma_{od}^2 & 0 \\ 0 & \sigma_{fl}^2 \end{pmatrix} \tag{M.9}$$

Although most noise is presumably caused by inhomogeneities in the culture liquid (e.g., through precipitation), mutual independence was assumed as OD and fluorescence were measured by different sensors over different media volumes and at different time instances.

#### S1.3 State Estimation

An EKF was employed that computed the state estimate with unimodal distribution. Linearization around the current estimate did not guarantee optimality, however, smooth system dynamics promised near optimality with the available models and measurements. The EKF computes the estimate in two steps: the prediction step and the measurement update step.

##### S1.3.1 Prediction Step

In every cycle (i.e., every minute), the EKF first predicted the current state based on the estimate from the last cycle and the production models derived in Equations (M.2) and (M.3). To minimize discretization errors, $t_s = 1$ s was chosen. Consequently, the prediction step ran 60 times each cycle. The dilution models in Equation (M.6) were added to the prediction when the bioreactor was diluting media to maintain the desired OD (typically set to 0.5).

##### S1.3.2 Measurement Update Step

Every last cycle of the growth period $k_h$, the measurement update step was augmented by the intermediate estimate $\hat{P}_{od}[k_h]$, $\hat{E}_{od}[k_h]$, and $\hat{P}_{fl}[k_h]$ that were derived from the measurements in the prior growth period.

The estimates through OD curvature (i.e., total growth rate) $\hat{P}_{od}[k_h]$ and $\hat{E}_{od}[k_h]$ were obtained by inserting the solution of Equation (M.1) into Equation (M.7):

$$\begin{aligned} od[k] &= P[k_h]e^{\mu_P(t[k]-t[k_h])} + E[k_h]e^{\mu_E(t[k]-t[k_h])} \\ &= P[k_h]e^{\mu_P(t[k]-t[k_h])} + (od[k_h] - P[k_h])e^{\mu_E(t[k]-t[k_h])} \end{aligned} \tag{M.10}$$

Rearranging Equation (M.10) and gathering each measurement from the first cycle $k_l$ to the last cycle $k_h$ of the growth period into vectors, resulted in:

$$\begin{pmatrix} e^{\mu_P(t[k_l]-t[k_h])} - e^{\mu_E(t[k_l]-t[k_h])} \\ \vdots \\ e^{\mu_P(t[k_h]-t[k_h])} - e^{\mu_E(t[k_h]-t[k_h])} \end{pmatrix} P[k_h] = \begin{pmatrix} od[k_l] - od[k_h]e^{\mu_E(t[k_l]-t[k_h])} \\ \vdots \\ od[k_h] - od[k_h]e^{\mu_E(t[k_h]-t[k_h])} \end{pmatrix} \tag{M.11}$$

In the matrix representation $\mathbf{A}P[k_h] = \mathbf{b}$, the least squares method can be applied to obtain $\hat{P}_{od}[k_h]$ that minimizes the residual in Equation (M.11). Finally, the algebraic relation in Equation (M.7) provides $\hat{P}_{od}[k_h]$.

The *P. putida* estimate $\hat{P}_{fl}[k_h]$ obtained through the curvature of the measured fluorescence was derived from the continuous model of Equation (M.3) and the solution of Equation (M.1):

$$\dot{F}(t) = \mu_F(t)P(t) = \mu_F P_h e^{\mu_P(t-t_h)} \tag{M.12}$$

Integration from all cycles within the growth period to cycle $k_h$ and rearrangement yielded:

$$\begin{pmatrix} \frac{\mu_F}{\mu_P}(e^{\mu_P(t[k_l]-t[k_h])} - 1) \\ \vdots \\ \frac{\mu_F}{\mu_P}(e^{\mu_P(t[k_h]-t[k_h])} - 1) \end{pmatrix} P[k_h] = \begin{pmatrix} F[k_l] - F[k_h] \\ \vdots \\ F[k_h] - F[k_h] \end{pmatrix} \tag{M.13}$$

Applying least-squares minimisation gave another *P. putida* estimate $\hat{P}_{fl}[k_h]$.

Having a good approximation of each method's exactness is vital when combining them. If the individual measurement updates are conflicting, the EKF should prioritize the more accurate one to derive a correct composition estimate. This was reflected in the EKF that adapted the influence of the measurement on the final composition estimate by altering its corresponding variance. During an experiment, the variance of the curvature methods was augmented by the least square fit, i.e., the normalized least-squares residual $SS_{norm}$:

$$SS_{norm} = \frac{SS_{res}}{SS_{tot}} \tag{M.14}$$

$SS_{res}$ is the residual sum of squares obtained from the least squares and $SS_{tot}$ is the total sum of squares, that is proportional to the variance of the data. The further $SS_{norm}$ was away from 0 and closer to 1 the worse was the least-square fit and the less trusted were intermediate estimates $\hat{P}_{od}[k_h]$, $\hat{E}_{od}[k_h]$ and $\hat{P}_{fl}[k_h]$. This is based on the correlation observed between the $SS_{norm}$ and the estimation error (Fig M1.2B). This metric of confidence in the estimates captured both estimation errors through temperature change, as well as those arising due to measurement noise, as both affected the quality of the least squares fit. The remaining quality limitations were not captured by the residual and thus were separately taken into account by increasing the individual uncertainty at the lower and higher temperature range or around the critical temperature. The uncertainties of the five measurements and estimates are reflected in the resulting time-varying measurement uncertainty matrix $R[k]$ (subsection S1.6).

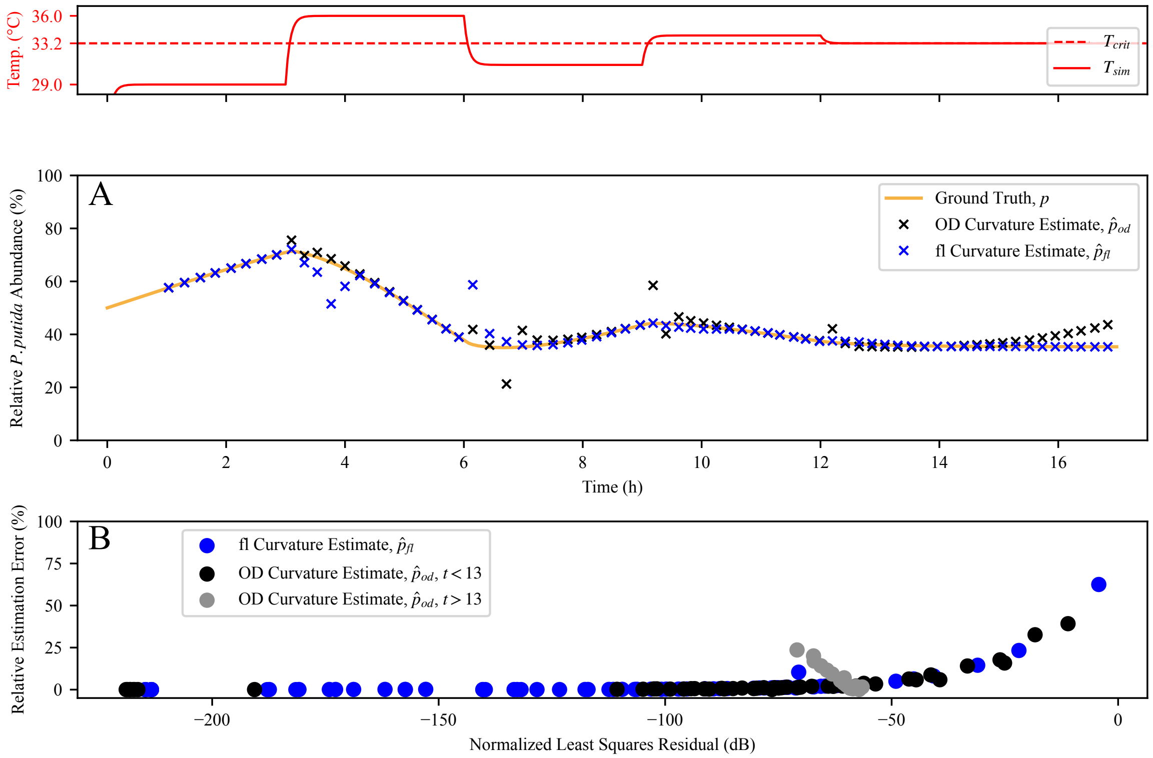


Figure M1.2: Noise-free simulation of the co-culture. Relative *P. putida* abundance estimation based on OD curvature $\hat{p}_{od}$ or fluorescence curvature $\hat{p}_{fl}$ over time and changing temperatures (A). The relative error of the different estimations over their normalized residual from the least squares fit (B). The data $\hat{p}_{od}$ is divided into situations before/after reaching the critical temperature around 13 h where growth-based estimation breaks down.

#### S1.4 Determining Bacteria Growth Rates

The derivation of the bacteria growth rates in the following is exemplified with *P. putida* but can be performed equivalently with *E. coli*. The growth rates were determined by growing the bacteria in monocultures, such that the OD measurements could be directly used to track bacteria abundance, e.g. $od[k] = P[k]$.

Between dilutions, i.e. in the growth period, the solution of the ODE in Equation (M.1) equals:

$$P(t) = P(t_l)e^{\mu_P(t-t_l)} \tag{M.15}$$

where $t_l$ corresponded to the point of time right after a dilution period. In the short time frame between dilutions, the growth rate $\mu_P$ was assumed to be constant. Instead of fitting $\mu_P$ to the exponential in Equation (M.15), the latter is transformed to obtain a first-order polynomial. Taking the natural logarithm and discretization leads to:

$$\begin{aligned} \ln P[k] &= \mu_P(t[k] - t[k_l]) + \ln P[k_l] \\ &= \mu_P t[k] + \ln P[k_l] - \mu_P t[k_l] \\ &= \mu_P t[k] + c \end{aligned} \tag{M.16}$$

In the next step a line can be fitted through all data points $k$ in the growth period. To account for the transformation and to obtain the optimal least-squares fit for the exponential function, the data points were weighted with $P[k]$. Finally, the growth rate was acquired by reading out the gradient of the fitted line.

#### S1.5 Determining Pyoverdine Production Rates

The Pyoverdine production rate $\mu_F$ can be determined by rearranging Equation (M.13):

$$\begin{pmatrix} \frac{P[k_h]}{\mu_P}(e^{\mu_P(t[k_l]-t[k_h])}-1) \\ \vdots \\ \frac{P[k_h]}{\mu_P}(e^{\mu_P(t[k_h]-t[k_h])}-1) \end{pmatrix} \mu_F = \begin{pmatrix} F[k_l]-F[k_h] \\ \vdots \\ F[k_h]-F[k_h] \end{pmatrix} \tag{M.17}$$

Where $\mu_P$ and $P[k_h]$ were obtained for each growth period as in subsection S1.4 and growth and production rates were assumed to be constant for the short time between dilutions. Applying an unweighted least-squares minimization gives the production rate $\mu_F$ that optimally fits the observed data in the growth period.

#### S1.6 Time-Varying Measurement Uncertainty Matrix

In the measurement update step, five different updates were formulated (subsubsection S1.3.2). This includes the measurements $od$ and $fl$ as well as the intermediate estimates $\hat{P}_{od}$, $\hat{E}_{od}$, and $\hat{P}_{fl}$. These can be compiled in the measurement vector $\mathbf{y}_{full}[k]$:

$$\mathbf{y}_{full}[k] = \begin{pmatrix} od[k] & fl[k] & \hat{P}_{od}[k] & \hat{E}_{od}[k] & \hat{P}_{fl}[k] \end{pmatrix}^T \tag{M.18}$$

The measurement uncertainty of $\mathbf{y}_{full}$ was then summarized in the matrix $\mathbf{R}_{full}[k]$ that got utilized in the measurement update step of the EKF:

$$\begin{aligned} \mathbf{R}_{full}[k] = diag(&\sigma_{od}, \\ &f_1(\sigma_{fl}, T[k]), \\ &f_2(\sigma_{P,od}, \mu_P[k], \mu_E[k], SS_{norm,od}[k]), \\ &f_2(\sigma_{E,od}, \mu_P[k], \mu_E[k], SS_{norm,od}[k]), \\ &f_3(\sigma_{P,fl}, T[k], SS_{norm,fl}[k])) \end{aligned} \tag{M.19}$$

Note that during cycles where the intermediate estimates are not available, $\mathbf{y}_{full}$ and $\mathbf{R}_{full}$ are reduced to:

$$\mathbf{y}[k] = \begin{pmatrix} od[k] & fl[k] \end{pmatrix}^T \qquad \mathbf{R}[k] = diag(\sigma_{od}, f_1(\sigma_{fl}, T[k]) \tag{M.20}$$

### References


[1] Steel H, Habgood R, Kelly CL, Papachristodoulou A. In situ characterisation and manipulation of biological systems with Chi.Bio. PLOS Biology. 2020;18(7):e3000794. doi:10.1371/journal.pbio.3000794.
